# Between fishing and farming: palaeogenomic analyses reveal cross-cultural interactions triggered by the arrival of the Neolithic in the Danube Gorges

**DOI:** 10.1101/2022.06.24.497512

**Authors:** Zuzana Hofmanová, Carlos S. Reyna-Blanco, Camille de Becdelièvre, Ilektra Schulz, Jens Blöcher, Jelena Jovanović, Laura Winkelbach, Sylwia M. Figarska, Anna Schulz, Marko Porčić, Petr Květina, Alexandros Tsoupas, Mathias Currat, Alexandra Buzhilova, Fokke Gerritsen, Necmi Karul, George McGlynn, Jörg Orschiedt, Rana Özbal, Joris Peters, Bogdan Ridush, Thomas Terberger, Maria Teschler-Nicola, Gunita Zariņa, Andrea Zeeb-Lanz, Sofija Stefanović, Joachim Burger, Daniel Wegmann

## Abstract

While early Neolithic populations in Europe were largely descended from early Aegean farmers, there is also evidence of episodic gene flow from local Mesolithic hunter-gatherers into early Neolithic communities. Exactly how and where this occurred is still unknown. Here we report direct evidence for admixture between the two groups at the Danube Gorges in Serbia. Analysis of palaeogenomes recovered from skeletons revealed that second-generation mixed individuals were buried amidst individuals whose ancestry was either exclusively Aegean Neolithic or exclusively local Mesolithic. The mixed ancestry is also reflected in a corresponding mosaic of grave goods. With its deep sequence of occupation and its unique dwellings that suggest at least semi-sedentary occupation since the late Mesolithic, the area of the Danube Gorges has been at the center of the debate about the contribution of Mesolithic societies to the Neolithisation of Europe. As suggested by our data, which were processed exclusively with uncertainty-aware bioinformatic tools, it may have been precisely in such contexts that close interactions between these societies were established, and Mesolithic ancestry and cultural elements were assimilated.

## Introduction

The Danube Gorges area, or Đerdap in Serbian, is situated in the Central Balkans and an early contact zone between migrating early European farmers and local foragers (Oross and Bánffy, 2009). With more than 25 prehistoric sites (Fig. 1), numerous Mesolithic and Neolithic human burial finds and a deep Final Paleolithic to Neolithic sequence of occupation it has contributed significantly to our understanding of the introduction of agriculture to the European continent and Mesolithic-Neolithic interactions (Porčić, Blagojević and Stefanović, 2016; de Becdelièvre *et al*., 2020).

**Figure 1.**
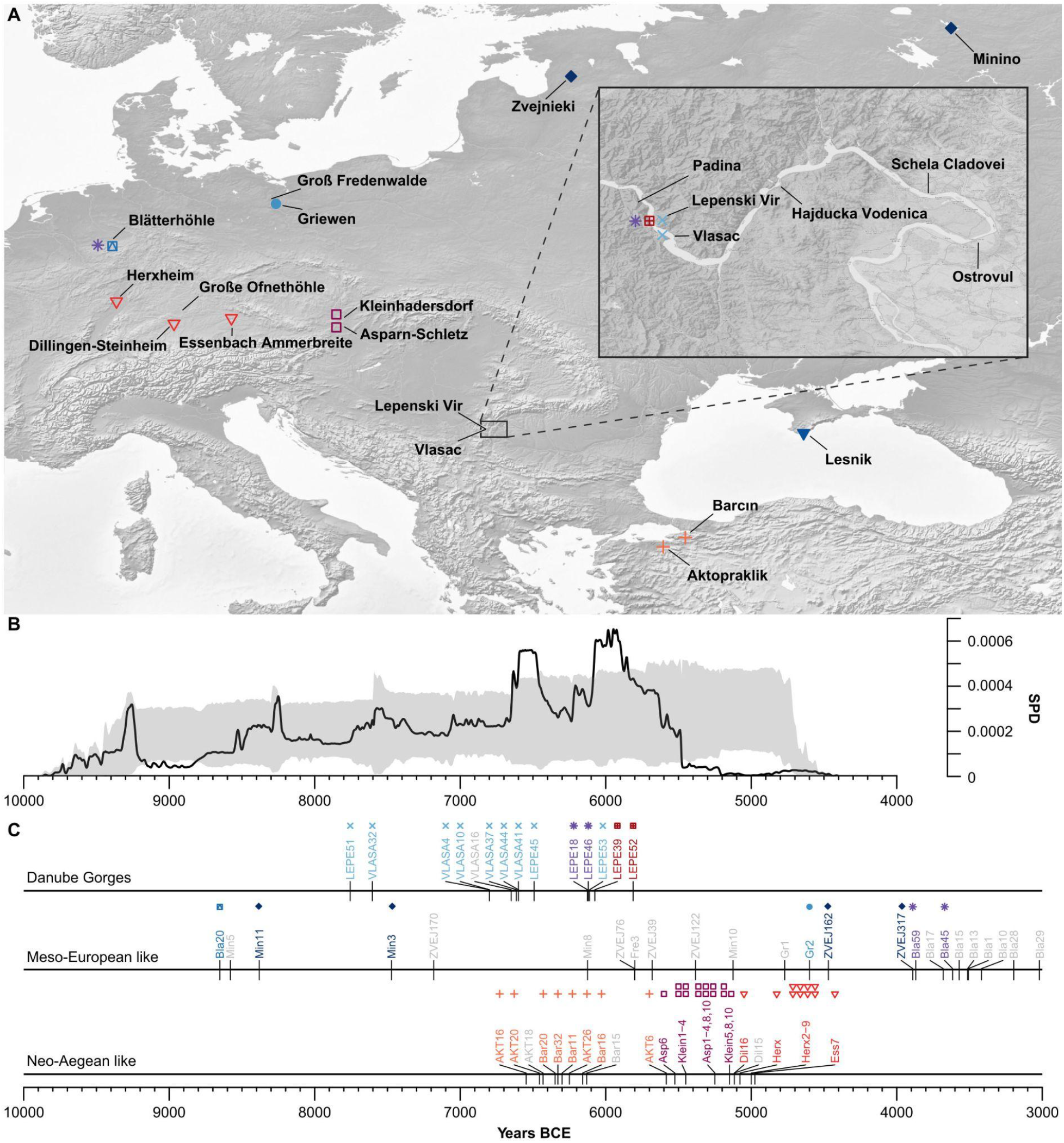
Spatial and temporal distribution of the ancient nuclear genomic samples sequenced in this study. **(A)** Location of archeological sites of Danube Gorges, North Aegean and Central, North and East Europe. **(B)** Observed summed probability distribution (SPD, solid black line) of calibrated radiocarbon dates from Danube Gorges along with a simulation envelope of the fitted model (shaded gray area). Adapted after (de Becdelièvre *et al*., 2021). **(C)** Chronological distribution of whole-genome and nuclear capture data sequenced in this study (see details in Table 1). Both symbols and labels are shown for each directly-14C or approximately dated sample, except for the ones who were filtered out for downstream analysis (labels shown in gray) or whose dates were not available.

**Table 1.** Archaeological and genetic information of all sequencing data either produced in this study or available for the Danube Gorges.

| Sample ID | Site | Phase<sup>a</sup> | Archaeological/<br>Burial ID | Age<br>(cal. BC) | Mean<br>Depth | Genetic<br>sex | R<sub>y</sub> | MT-Haplogroup |
| --- | --- | --- | --- | --- | --- | --- | --- | --- |
| **Danube Gorges whole genomes** |  |  |  |  |  |  |  |  |
| VLASA16* | Vlasac | LM | VL 53 | 6,650 ± 174 | 0.99 | XX | 0.0012 | U5a1c |
| VLASA37 | Vlasac | LM | VL 24 | 6,614 ± 153 | 5.06 | XY | 0.0829 | K1f |
| LEPE45 | Lepenski Vir | TEN | LV 91 | 6,492 ± 96 | 4.03 | XY | 0.0754 | U5a2d |
| LEPE46*.<sup>#</sup> | Lepenski Vir | TEN | LV 93 | 6,120 ± 102 | 0.74 | XX | 0.0014 | H |
| LEPE51 | Lepenski Vir | EM | LV 68 | 7,756 ± 184 | 3.74 | XX | 0.0004 | U4a2 |
| LEPE53*.<sup>#</sup> | Lepenski Vir | TEN | LV 27/a | 6,111.5 ± 104.5 | 0.91 | XX | 0.0013 | U5b2c1 |
| **Danube Gorges neutralomes** |  |  |  |  |  |  |  |  |
| VLASA4 | Vlasac | LM | VL 18/a | ~7,400-6,200 | 85.82 | XY | 0.0815 | U5a1c |
| VLASA10 | Vlasac | LM | VL 41 | ~7,400-6,200 | 41.74 | XY | 0.0978 | K1f |
| VLASA32<sup>#</sup> | Vlasac | LM | VL 16 | 7,604.5 ± 136.5 | 43.08 | XY | 0.08 | U5a2a |
| VLASA41 | Vlasac | LM | VL 30 | ~7,400-6,200 | 55.97 | XX | 0.0005 | U5b2b |
| VLASA44 | Vlasac | LM | VL 47 | ~7,400-6,200 | 64.96 | XY | 0.0819 | U5b2b |
| LEPE18 | Lepenski Vir | TEN | LV 27/d | 6,126 ± 100 | 78.71 | XY | 0.0822 | U5a2 |
| LEPE39 | Lepenski Vir | TEN | LV 82 | 6,075 ± 125 | 23.26 | XY | 0.0798 | T2e |
| LEPE46<sup>#</sup> | Lepenski Vir | TEN | LV 93 | 6,120 ± 102 | 52.09 | XX | 0.0005 | H |
| LEPE52<sup>#</sup> | Lepenski Vir | E-MN | LV 73 | 5,812 ± 119 | 61.44 | XY | 0.082 | H2a |
| LEPE53<sup>#</sup> | Lepenski Vir | TEN | LV 27/a | 6,111.5 ± 104.5 | 36.39 | XX | 0.0004 | U5b2c1 |
| **Danube Gorges whole genomes - published data (Marchi *et al.*, 2022)** |  |  |  |  |  |  |  |  |
| VLASA7 | Vlasac | LM | VL 31 | 6,552 ± 212 | 15.21 | XY | 0.0754 | U5a2a |
| VLASA32<sup>#</sup> | Vlasac | LM | VL 16 | 7,604.5 ± 136.5 | 12.66 | XY | 0.0753 | U5a2a |
| LEPE48 | Lepenski Vir | TEN | LV 122 | 5,939.5 ± 72.5 | 10.93 | XY | 0.0755 | K1a1 |
| LEPE52<sup>#</sup> | Lepenski Vir | E-MN | LV 73 | 5,812 ± 119 | 12.38 | XY | 0.0751 | H3 |
| **non-Danube Gorges nuclear capture genomes** |  |  |  |  |  |  |  |  |
| Lec2 | Lesnika Cave,<br>Crimea Ai-Petri | EM | Lsa 031 | - | 12.906 | XY | 0.0952 | U5a1 |
| Min2 | Minino | M | Minino I, 11 | - | 16.212 | XX | 0.0006 | U4 |
| Min3 | Minino | M | Minino II, III | 7,472 ± 52 | 58.044 | XY | 0.089 | U4a1 |
| Min5* | Minino | M | Minino I, 3 | 8,580 ± 160 | 3.365 | XY | 0.077 | U4 |
| Min8* | Minino | M | Minino I, 5 | 6,125 ± 325 | 0.812 | XY | 0.0826 | U4a1 |
| Min10* | Minino | M | Minino I, 20 | 5,125 ± 525 | 3.917 | XX | 0.0006 | U4d |
| Min11 | Minino | M | Minino II, V | 8,671 ± 48<br>8,092 ± 94 | 39.340 | XY | 0.1181 | U4a1 |
| ZVEJ317 | Zvejnieki | M | ZV317 | 3,890 ± 67 | 28.716 | XY | 0.1415 | U4a1 |
| ZVEJ39* | Zvejnieki | M | ZV39 | 5,681 ± 36 | 1.707 | XY | 0.0788 | U5a2c |
| ZVEJ76* | Zvejnieki | M | ZV76 | 5,802 ± 73 | 1.164 | XX | 0.0006 | U5a2 |
| ZVEJ122* | Zvejnieki | M | ZV122 | 5,383 ± 68 | 0.971 | XY | 0.0889 | U5a2d |
| ZVEJ162 | Zvejnieki | M | ZV162 | 4,470 ± 72 | 17.264 | XY | 0.1132 | U4a1 |
| ZVEJ170* | Zvejnieki | M | ZV170 | 7,182 ± 107 | 0.251 | XY | 0.0793 | U5a2d |
| Bla20 | Blätterhöhle | M | BH 04/174 | 8,652 ± 58 | 6.558 | XX | 0.0005 | U5a2c |
| Fre3* | Groß Fredenwalde | M | Ind 1 | 5,800 ± 400 | 0.315 | XY | 0.0762 | H2a2a |
| Gr1* | Criewen | M | Criewen<br>1961:82 Grab1 | 4,770 ± 40 | 5.104 | XY | 0.1072 | U5b2a2 |
| Gr2 | Criewen | M | 1961: 82/7 | 4,600 ± 60 | 21.134 | XX | 0.0005 | U4b1b1 |
| GrO1* | Große Ofnethöhle | LM | Ind 12, Ktg nr<br>SV 001/12<br>(2485) | - | 11.351 | XY | 0.0795 | U5b1d1 |
| AKT6 | Aktopraklik | EN | H17/50.1 | 5,584 ± 49 | 32.182 | XY | 0.0815 | T2 |
| AKT16# | Aktopraklik | EN | D89/14.1 | 6,547 ± 87 | 39.813 | XX | 0.0019 | K1a3 |
| AKT18* | Aktopraklik | EN | D89/17.1 | 6,431 ± 38 | 3.76 | XX | 0.0006 | H2a2a |
| AKT20 | Aktopraklik | EN | E89/9.3 | 6,456 ± 37 | 20.96 | XY | 0.0775 | J2b1 |
| AKT26 | Aktopraklik | EN | D90/4.4 | ~6,500 - 6,000 | 18.77 | XX | 0.0004 | J2a |
| Bar11 | Barcin | EN | M11/93 | ~6,600 - 6,000 | 24.359 | XY | 0.0801 | X |
| Bar15*.<sup>1</sup> | Barcin | EN | M10/115 | 6,131 ± 82 | 25.558 | XX | 0.0004 | K1a2 |
| Bar16 | Barcin | EN | L10/187 | 6,159 ± 74 | 22.579 | XX | 0.0004 | K1a1 |
| Bar20 | Barcin | EN | M11S/401 | 6,348 ± 90 | 13.561 | XX | 0.0006 | W5 |
| Bar32 | Barcin | EN | L11/604 | 6,329 ± 90 | 13.688 | XX | 0.002 | K1a2 |
| Asp1 | Asparn-Schletz | EN (LBK) | 93/7//93/8, Nr.<br>4451/Schnitt<br>22 | 5,250 ± 250 | 21.639 | XY | 0.0945 | K1a |
| Asp2 | Asparn-Schletz | EN (LBK) | Ind 10, S6/<br>282 | 5,250 ± 250 | 14.771 | XY | 0.085 | N1a1a1a2 |
| Asp3 | Asparn-Schletz | EN (LBK) | Nr.<br>4455/Schnitt<br>22 1993/19 | 5,250 ± 250 | 33.465 | XX | 0.0018 | J1c17 |
| Asp4 | Asparn-Schletz | EN (LBK) | 2490, Ind 7,<br>7//restl.<br>Schädel 5/02 | 5,250 ± 250 | 29.061 | XX | 0.0021 | T2b13b |
| Asp6# | Asparn-Schletz | EN (LBK) | Ind 44 (646<br>Part 152,<br>Schnitt 10<br>LM70 | 5,525 ± 50 | 23.815 | XY | 0.0928 | U5a1c1 |
| Asp8 | Asparn-Schletz | EN (LBK) | Ind 24, 374<br>S7/LM72 Grab<br>1 | 5,250 ± 250 | 76.578 | XX | 0.0019 | X2b |
| Asp10 | Asparn-Schletz | EN (LBK) | Ind 6 | 5,250 ± 250 | 48.450 | XY | 0.1047 | K1a |
| Klein1 | Kleinhadersdorf | EN (LBK) | 25,939 | ~5,950 - 5,150 | 85.4511 | XX | 0.0018 | T2b |
| Klein2 | Kleinhadersdorf | EN (LBK) | 25,941 | ~5,950 - 5,150 | 44.729 | XX | 0.0019 | H |
| Klein3 | Kleinhadersdorf | EN (LBK) | 25,945 | ~5,950 - 5,150 | 30.345 | XX | 0.002 | J1c2 |
| Klein4 | Kleinhadersdorf | EN (LBK) | 25,926 | ~5,950 - 5,150 | 49.749 | XX | 0.0019 | T2b |
| Klein5 | Kleinhadersdorf | EN (LBK) | 25,925 | ~5,950 - 5,150 | 44.777 | XY | 0.0957 | N1a1a1a3 |
| Klein8 | Kleinhadersdorf | EN (LBK) | 25,923 | ~5,950 - 5,150 | 41.197 | XX | 0.0006 | U5b |
| Klein10 | Kleinhadersdorf | EN (LBK) | 25,936 | ~5,950 - 5,150 | 49.393 | XX | 0.0021 | N1a1a1 |
| Dil15*.<sup>1</sup> | Dillingen-Steinheim | EN (LBK) | Grab 23,<br>Befund 24 | 5,116 ± 118 | 28.668 | XY | 0.0918 | J1c |
| Dil16#.<sup>1</sup> | Dillingen-Steinheim | EN (LBK) | Grave 24,<br>Befund 24 | 5,116 ± 118 | 25.186 | XY | 0.0952 | J1c |
| Ess7# | EssenbachAmmerbreit | EN (LBK) | Grave 2 | 4,975 ± 75 | 12.2172 | XY | 0.0793 | U5b2c1 |
| Herx# | Herxheim | EN (LBK) | 281-19-6 | 5,078 ± 85 | 9.23 | XX | 0.0004 | K1a4a1i |
| Herx2 | Herxheim | EN (LBK) | 282-126-7 | ~5,000 | 55.87 | XY | 0.1266 | J2b1 |
| Herx3 | Herxheim | EN (LBK) | 282-13-7 | ~5,000 | 16.14 | XX | 0.0006 | W1+119 |
| Herx4 | Herxheim | EN (LBK) | 282-23-1 | ~5,000 | 115.12 | XY | 0.1039 | K1a |
| Herx5 | Herxheim | EN (LBK) | 282-94-11 | ~5,000 | 30.32 | XY | 0.087 | HV+16311 |
| Herx6 | Herxheim | EN (LBK) | 7034-12 | ~5,000 | 36.42 | XX | 0.0005 | U3 |
| Herx7 | Herxheim | EN (LBK) | 282-104-4 | ~5,000 | 34.39 | XY | 0.0787 | K1a1a |
| Herx8 | Herxheim | EN (LBK) | 282-126-16 | ~5,000 | 28.77 | XX | 0.0009 | T2e |
| Herx9 | Herxheim | EN (LBK) | 282-88-2 | ~5,000 | 57.05 | XX | 0.0006 | K1a+150 |
| Bla1* | Blätterhöhle | MN | BH 04/011 | 3,508 ± 102 | 0.689 | XY | 0.0981 | H1bm |
| Bla10* | Blätterhöhle | MN | BH 04/012 B | 3,418 ± 63 | 3.528 | XY | 0.0781 | H2a2a |
| Bla13* | Blätterhöhle | MN | BH 04/034 | 3,513 ± 102 | 4.549 | XX | 0.0006 | H2a2a |
| Bla15* | Blätterhöhle | MN | BH 04/041 | 3,571 ± 47 | 0.509 | XY | 0.0845 | H2a2a1 |
| Bla17* | Blätterhöhle | MN | BH 04/032 | 3,681 ± 19 | 1.663 | XY | 0.1036 | H1ba |
| Bla28* | Blätterhöhle | MN | BV 06G5d/Po. 49.2 | 3,196 ± 103 | 0.4235 | XY | 0.0813 | J1c1 |
| Bla29* | Blätterhöhle | MN | BH 07 I4/Po 1 | 3,020 ± 61 | 1.282 | XX | 0.0009 | H2a2a1 |
| Bla75* | Blätterhöhle | MN | BH14<br>Qu7c/Bla75 | - | 0.303 | ~XY | 0.3047 | - |
| Bla32 | Blätterhöhle | MN | BH 04/072 | - | 47.839 | XX | 0.0005 | H5 |
| Bla45 | Blätterhöhle | MN | BH14<br>Qu7c/Bla45 | 3,616 ± 56<br>3,922 ± 60 | 32.4236 | XY | 0.095 | U5b2b |
| Bla59 | Blätterhöhle | MN | BH14<br>Qu7c/Bla59 | 3,869 ± 59 | 10.879 | XY | 0.0943 | U5b2a2 |

<sup>a</sup> M, Mesolithic; EM, Early Mesolithic; LM, Late Mesolithic; TEN, Transformation/Early Neolithic; EN, Early Neolithic; E-MN, Early-Middle Neolithic; MN, Middle Neolithic.
\* Samples were discarded for downstream analysis.

After a Final Paleolithic occupation in some rock-shelter sites (ca. 13,000-9,500 cal BC), the human presence at the Danube Gorges is documented on open-air sites in river terraces at several locations, such as Lepenski Vir and Padina during the Early Mesolithic period (ca. 9,500-7,400/7,300 cal BC; (Boroneanţ, 1999; Borić, 2011)). The population then likely became more numerous during the Late Mesolithic period (ca. 7400/7300-6300/6200 cal BC), and some finds indicate that the Mesolithic population lived partly semi-sedentary lives and their mobility was greatly reduced compared to earlier periods (Dimitrijević, Živaljević and Stefanović, 2016; de Becdelièvre *et al*., 2021). This semi-sedentary lifestyle was undoubtedly associated with the consumption of aquatic food: the community relied heavily on fishing and benefited especially from the large numbers of anadromous species that populated the narrow gorge during their upward migration (Bonsall *et al*., 1997; Bartosiewicz *et al*., 2001; Nehlich *et al*., 2010; Borić *et al*., 2014; Jovanović *et al*., 2019). The diet was further enriched by game, different plant species, and possibly dogs (Cristiani *et al*., 2016; Jovanović *et al*., 2019, 2021).

At the end of the 7th millennium BC, the first settlements with farming subsistence appeared in the Central Balkans and the southern part of the Pannonian plain: the Neolithic Starčevo-Körös-Criş cultural complex (Garašanin, 1982; Tringham, 2000; Krauß, 2011), which later developed important sites, such as Starčevo and the early layers of Vinča tell, in the immediate vicinity of the Danube Gorges (Porčić, Blagojević and Stefanović, 2016; Jovanović et al., 2019; de Becdelièvre et al., 2020; Porčić et al., 2020, 2021). Around this time, trapezoidal houses were built at the Lepenski Vir site. Such houses, while similar to Late Mesolithic dwellings at the nearby site Vlasac, have a shape not known outside the Danube Gorges (Boric, French and Dimitrijević, 2008), but might have some structural analogies, such as plastered floors, in Neolithic Anatolia (Srejović, 1969; Borić, 2011; Borić, Radović and Stefanović, 2012). Mesolithic foragers and newcomers with a Neolithic background used the settlement site during this period simultaneously (Borić and Price, 2013; Hofmanová, 2016; González-Fortes et al., 2017; Mathieson et al., 2018) and, as indicated by isotopic data in skeletons from the time of their arrival, newcomers had a more terrestrial diet (Borić and Price, 2013). This period of interaction between foragers and farmers at the site is referred to as the “Transformation Phase” and dates ca 6200-5950 cal BC (Borić et al., 2018). Besides the trapezoidal houses, other elements of the Neolithic way of life, such as typical ornaments, raw materials and ceramics, now appear (Garašanin and Radovanović, 2001). In addition, so-called ancestor statues with fish-like features have been found in the corresponding layers of Lepenski Vir. Over time, Neolithic features increased at the site, leading to the fully developed Neolithic period (5,950-5,550 cal BC), in which the trapezoidal houses were abandoned (citation), the Neolithic suite of domesticated animals appeared (Boric and Dimitrijevic, 2005; Borić and Dimitrijević, 2007) and grains of cereals have been evidenced in the dental calculus of some Neolithic individuals (Jovanović et al., 2021). However, the use of wild resources remained dominant (Boric and Dimitrijevic, 2005; Borić and Dimitrijević, 2007; Cramp et al., 2019; Jovanović et al., 2019).

This wealth of evidence makes Lepenski Vir one of the best-studied sites of the Mesolithic-Neolithic transition and provides a unique opportunity to explore foragers and farmers interactions. However, key aspects of the site’s population history remain unclear, including how the mixing between the Danubian and Aegean Neolithic members of the society took place, i.e. when and with what proportions of the two populations it took place. Furthermore, it is not known how these findings from the Danubian Gorges relate to other forager or farming populations in SW Asia and Europe.

Using uncertainty-aware bioinformatic tools on high quality genomic data from 51 available and 50 newly produced samples with a geographical and chronological focus on the Meso-Neolithic transition in the Danube Gorges and key sites from the Central Balkans, Lower Austria, Southern Germany, the Marmara region and the Baltics, we show that the two contributing populations were genetically well differentiated but episodically interacted, as evidenced not least by the presence of first generation mixed individuals.

## Results

In order to better understand the reciprocal relationship between the populations originating from the Neolithic area in the wider Aegean including the Sea of Marmara and the indigenous foragers in Serbia, we produced (Table 1): 1) six whole genomes at a sequencing depth of 1-5X and ten neutralomes of 5Mb length (see Methods) from the Danube Gorges, 2) ten Neolithic neutralomes from the Aegean/Marmara region 3) 37 neutralomes from Neolithic Central Europe, 4) five neutralomes from Mesolithic Central Europe, 6) 12 neutralomes from hunter-gatherers of North-Eastern Europe, 7) one neutralome from Lesnik Cave presumably dating to the Final Paleolithic, and 8) 78 mtDNA genomes from the same regions (Supplementary Data Table 1). We complemented these with 53 chronologically similar whole genomes available from Europe, NW and Central Anatolia and the Caucasus ((Gamba *et al*., 2014; Lazaridis *et al*., 2014; Olalde *et al*., 2014, 2015; Skoglund *et al*., 2014; Jones *et al*., 2015, 2017; Broushaki *et al*., 2016; Hofmanová *et al*., 2016; Kılınç *et al*., 2016; González-Fortes *et al*., 2017; Sikora *et al*., 2017; Günther *et al*., 2018; Marchi *et al*., 2022), Table S1), including four from the Danube Gorges ((Marchi *et al*., 2022), Table 1), and 20 modern samples from Africa and West Eurasia retrieved from the SGDP database ((Mallick *et al*., 2016), Table S1).

To ensure comparability and increase the sensitivity of population genomic analyses, we focused on 54 whole genomes and 54 5Mb neutralomes (with seven overlaps, Table 1, S1) that passed rigorous quality assessments mainly evaluating the reproducibility of genetic diversity estimates in face of data bootstrapping (see Methods, Figure S1-S2). Further, we based our analyses on uncertainty-aware inference methods that use genotype likelihoods, which were inferred from raw sequence data (unless not publicly available) using a bioinformatic pipeline dedicated to ancient DNA (see Methods, Supplementary Data Table 1; (Link *et al*., 2017)). Finally, we excluded an individual from Dillingen, Germany (Dil15), which we identified as a brother of Dil16, as well as one from Barcın, Turkey (Bar15), which we identified to have a parent-child relationship with Bar8 (Figure S2D).

### Early-generation admixed individuals at the Danube Gorges

Two Danube Gorges individuals, LEPE18 (LV 27d, 6,126 ± 100 cal BC) and LEPE46 (LV 93, 6,120 ± 102 cal BC), display substantial ancestry from both clusters. To shed more light on their admixture status, we used a Bayesian approach (Shastry *et al*., 2021) to infer genome-wide ancestry proportions (q_1_) jointly with inter-population ancestry proportions (Q_12_), i.e. the fraction of the genome at which a sample is heterozygous for the different ancestries (Fig 2D). These estimates indicate that both samples were second-generation admixed individuals: both had a mixed first-generation parent, while the other parent was unmixed of either Meso European-like (LEPE18) or Neo Aegean-like ancestry (LEPE46). Thus, LEPE46 had one Meso European-like and three Neo Aegean-like grandparents, while LEPE18 had one Neo Aegean-like and three Meso European-like grandparents.

**Figure 2.**
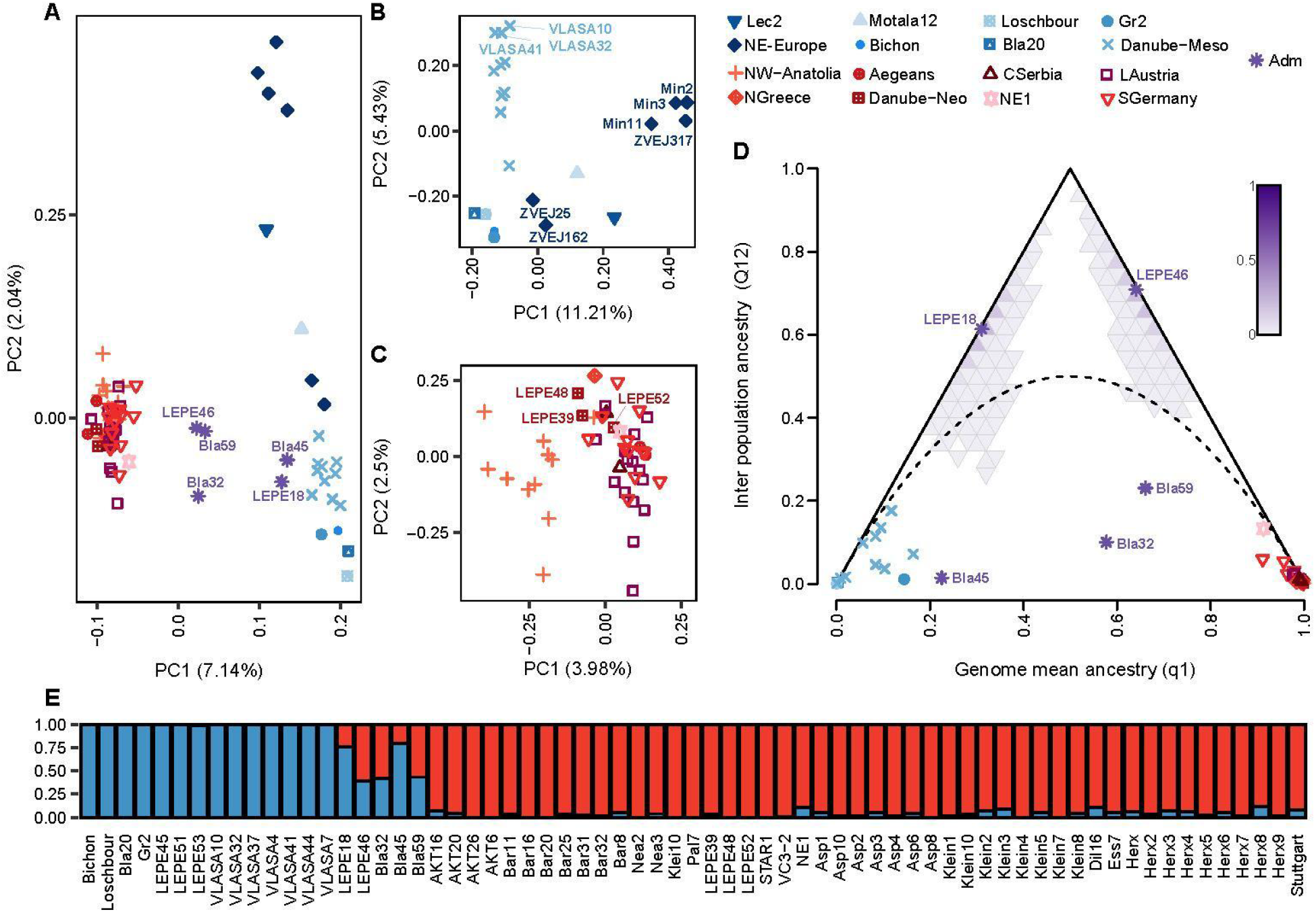
PCA and Admixture of Meso European-like and Neo Aegean-like individuals. (**A**) PCA that includes Neo Aegean-like individuals in red-like colors and Meso European-like individuals in blue-like colors. Admixed individuals are shown in purple. (**B**) PCA using only Meso European-like populations. (**C**) PCA using only Neo Aegean-like populations. (**D**) Entropy analysis with the same dataset as in (A) but excluding Lec2, Motala12 and North-Eastern European hunter-gatherers. The y axis denotes the proportion of the genome in which one gene copy comes from Neo Aegean-like ancestry and the other from Meso European-like ancestry; the position of our Lepenski Vir individuals are associated with backcross lineages. The x axis denotes the proportion for Neo Aegean-like genome ancestry estimated by Entropy. Inner triangles represent the samples of the posterior distribution and the curved dotted line is based on Hardy-Weinberg equilibrium (HWE). (**E**) NGSadmix results for K=2 using the set from Entropy (D); blue represents the Meso European-like ancestry and red the Neo Aegean-like ancestry.

We identified three additional individuals with considerable ancestry from both clusters: Bla32, Bla59 and Bla45 from the Blätterhöhle cave in Westfalia, Germany (Fig. 2A) dating to 4th millennium cal BC (3900-3000 cal BC). In contrast to LEPE18 and LEPE46, they appear to be later generation admixed (Fig. 2D), in line with their more recent age (see Table 1) and the previously reported ongoing admixture at this site (Bollongino *et al*., 2013; Lipson *et al*., 2017) and in the wider region (Haak *et al*., 2015).

### High genetic diversity in Danube Gorges foragers

To characterize the forager population present at the Danube Gorges during the Mesolithic and the Transformation phase, we first focused on individuals with <4% Neo Aegean-like ancestry in the admixture analysis. As attested by a projection-free PCA (Figure 2B), the Meso European-like individuals from the Danube Gorges are most similar to those from Western Europe, albeit slightly shifted towards individuals from North-Eastern Europe, in line with previous reports (Mathieson *et al*., 2018). In contrast to the Western European individuals, their cluster appears rather diverse (Figure 2B), which likely reflects a locally large population, elevated gene flow from neighboring populations, or both. This interpretation, consistent with the idea of a partially sedentary and prosperous fishing society of the Transformation period, is corroborated by Danube Gorges individuals generally having the highest genome-wide heterozygosity levels and shortest total lengths of runs of homozygosity (ROH) among all post-LGM Meso European-like individuals (Figure 3A, B, E), albeit some individual variation. Interestingly, the three Danube Gorges individuals from the Vlasac site that fall most distantly from the other Western European Meso European-like samples on the PCA (Fig 2A, B, Figure S3B, VLASA10, VLASA32, VLASA41) were among the only four buried with disarticulated skulls.

**Figure 3.**
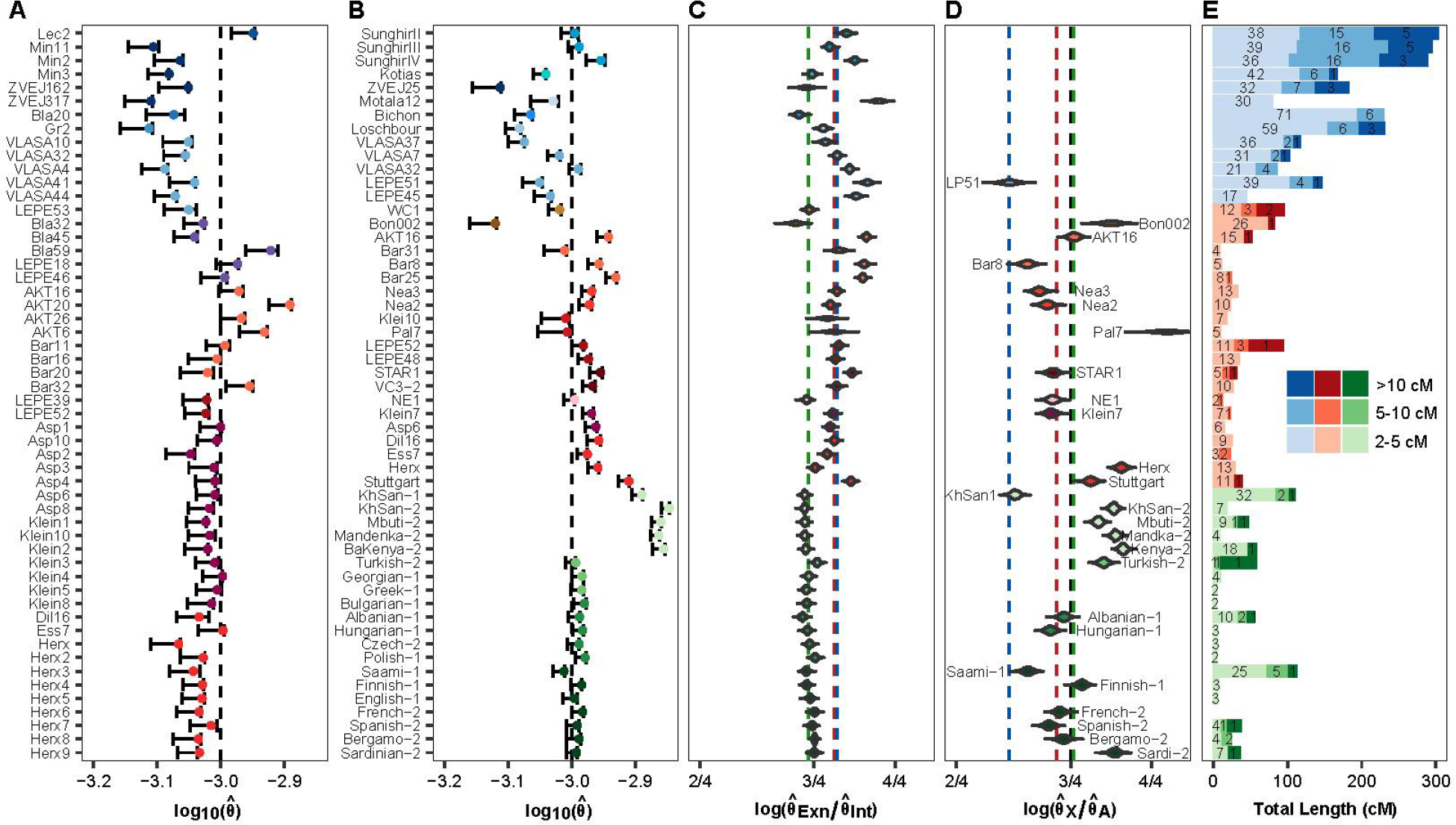
Intra and inter genetic diversity of ancient and modern samples. (**A**) Heterozygosity (*θ*) inferred genome-wide along the 5Mb neutral capture sites for neutralome samples. The dashed line indicates the expected value, the points represents the MLE estimate and the error bar the min/max of 100 bootstrap values. (**B**) Heterozygosity (*θ*) inferred genome-wide along the ∼18Mb customized neutral genome sites for whole-genome samples. (**C**) Exon/Intron heterozygosity (*θ*) ratio inferred for whole-genome samples. The dashed lines indicate the median of each group ratio value. (**D**) Heterozygosity (*θ*) ratio estimated between X neutral sites (∼11Mb) and autosomal neutral genome sites (∼18Mb) for whole-genome female samples. The black dashed line indicates the expected X/A ratio value (3/4), the other colored dashed lines represent the median of each group ratio value. (**E**) Runs of homozygosity (ROHs) for all whole-genome samples; the fill color represents the broad cultural periods, the genetic lengths were binned in 2-5 cM, 5-10 cM and >10 cM, and the observed counts are written inside each bar category.

A similarly diverse cluster is observed for the individuals of the site of Zvejnieki, with the two older samples ZVEJ25 (grave 93, 5,738 ± 102 cal BC) and ZVEJ162 (grave ZV162, 4,470 ± 72 cal BC) clustering with Meso European-like samples from Western Europe, while the youngest sample ZVEJ317 (grave ZV317, 3,890 ± 67 cal BC) does not (Figure 2B). The influx of a rather distinct ancestry into Zvejnieki during the Neolithic has been previously reported (Jones *et al*., 2017). As our data shows, the source of the influx was genetically very close to samples from Minino, which lies around 1,600 km to the east. Since the Baltic Sea region was covered with ice until at least the late 10th millennium cal BC, it is reasonable to assume that the two distinct ancestors discovered at Zvejnieki ultimately came from two different glacial refugia of the late Ice Age, possibly one in southern France and one on the Black Sea coast (Mittnik *et al*., 2018). Despite this diverse origins, we estimate low levels of heterozygosity for all Zvejnieki and Minino samples but no evidence of recent inbreeding for ZVEJ25 (the only sample with sufficient quality whole genome data).

Noteworthy, we estimated the lowest heterozygosity among all neutralomes for the late hunter-gatherer from the site Criewen (GR2, (Terberger *et al*., 2018)), a female from Northern-Eastern Germany, with a date around 4,500 cal BC (dated to 4,600 ± 60 cal BC but likely with reservoir effect). It represents the most recent Central European individual with essentially 100% Meso European-like ancestry analyzed here.

### Neolithic ancestry in Lepenski Vir resembles that from Early Neolithic Northern-Greece

To characterize the Neolithic population appearing at the Danube Gorges around 6,200 cal BC, we next focused on all samples with >96% Neo Aegean-like ancestry in the Admixture analysis. On a projection-free PCA (Figure 2C, Figure S3A), the first axis separates Neo Aegean-like samples of the Marmara region (Aktopraklık and Barcın sites) from those of Greece and Central Europe, a distinction usually not seen on projected PCAs (e.g. (Mathieson *et al*., 2018)). The three Neo Aegean-like individuals from Lepenski Vir as well as Greek individuals from the same time (Nea2, Nea3) fall closest to those from Marmara, unlike the Greek individuals from later periods (Pal7, Klei10). The same chronological signal is also seen among the Lepenski Vir individuals, among which the two from the Transformation Period, LEPE48 (LV 122, 5,939.5 ± 72.5 cal BC) and LEPE39 (LV 82, 6,075 ± 125 cal BC) appear closer to Marmara samples than LEPE52 (LV 73, 5,812 ± 119 cal BC), the sample from the later period of the fully developed Neolithic. This could suggest extended gene flow with other Neolithic sites in the later phases of Lepenski Vir, which would also explain the observed shift in the composition of the mtDNA gene pool towards those sites over time (Supplementary Data Table 1). However, as far as the further Neolithic expansion along the Balkan route from the Aegean to Central Europe is concerned, the PCA does not show a clear spatiotemporal signal of differentiation. This also applies to the individuals from Asparn Schletz and Herxheim, which are thought to derive from a massacre or ritual background potentially involving individuals from a larger area (Boulestin *et al*., 2009; Orschiedt and Haidle, 2012; Schulting and Fibiger, 2012; Boulestin, 2015). At the level of genomic depth studied here, they do not appear particularly diverse, comparable to the individuals from more regular Neolithic burial sites such as Kleinhadersdorf.

### Heterozygosity is higher in farmers than in foragers and decreases along the Neolithic route of expansion

With the exception of a single individual from the Herxheim site in SW-Germany (grave 281-19-6, 5,078 ± 85), the genome-wide heterozygosity estimated from neutralomes for all Neo Aegean-like individuals was consistently higher than for post-LGM Meso European-like individuals, both overall as well as when comparing samples from the Danube Gorges only (Fig 3A). In line with the idea of the Neolithic expansion along a Balkan route, we found generally the highest diversity both in terms of differentiation on the PCA (Figure 2C) and heterozygosity (Figure 3A,B) in the Marmara region (Aktopraklık and Barcın). While there is a general spatial trend of decreasing heterozygosity from the Marmara region towards SW Germany, there appears to be no temporal change in the Marmara region. Rather, the Chalcolithic sample AKT6 (grave H17/50.1; 5,584 ± 49 cal BC), which dates ∼1000 years later, shows heterozygosity comparable to that of the Early Neolithic skeletons from the same region.

Consistent with their admixed status, the genome-wide heterozygosity of LEPE18 and LEPE46 was estimated above that of most Meso European-like and Neo Aegean-like samples, especially those from the Danube Gorges.

Heterozygosity estimates from whole genomes confirmed these general patterns, albeit the smaller sample size. The heterozygosity of Neo Aegean-like samples appears to match that of modern Europeans (Fig 3B), with the exception of Bar31, Klei10 and Pal7 that had slightly lower levels similar to that of WC1 from Wezmeh Cave, an Early Neolithic individual from Iran with a very different demographic history (Broushaki *et al*., 2016; Marchi *et al*., 2022). For Bon002, we inferred particularly low levels of heterozygosity, in line with previous reports (Kılınç *et al*., 2016).

The lower heterozygosity of post-LGM foragers compared to Neolithic individuals was previously seen at larger geographic scales (Fu *et al*., 2016; Kılınç *et al*., 2016; Posth *et al*., 2016; Kousathanas *et al*., 2017; Renaud *et al*., 2019) and interpreted as a result of their long-term demography, such as a severe bottleneck during the LGM and their generally low population size (Gamble *et al*., 2004; Fernández-López de Pablo *et al*., 2019; Marchi *et al*., 2022). In support of a strong LGM bottleneck, we estimated elevated genome-wide diversity for the Meso European-like individuals from Sunghir predating the LGM (Sikora *et al*., 2017). In support of low post-LGM population size, we found all Meso European-like individuals, including those from the Danube Gorges, to have longer total length of runs of homozygosity (ROH) than Neolithic samples (but not LEPE52, which was likely recently inbred) (Fig 3E). Despite their elevated diversity, however, the three pre-LGM Sunghir individuals had the longest total ROH and the highest number of very long ROH segments of all Meso European-like individuals, in line with the interpretation of very small populations embedded in larger mating networks (Sikora *et al*., 2017). Interestingly, we estimated elevated diversity comparable to that of the Sunghir samples also for Lec2, an individual from Lesnik Cave with unclear dating but possible of pre-LGM origin.

### No evidence for strong purifying selection in the Neolithic population

We next evaluated whether these demographic events led to a reduction in the efficacy of purifying selection by inferring the heterozygosity at exons relative to that at introns for each whole-genome sample (Fig 3C). As expected, we estimated lower diversity at exons than at introns for all individuals, but found this reduction was much more pronounced in modern than ancient individuals, which hints at increased purifying selection in recent times, probably as a result of population growth (Gazave *et al*., 2013). However, we inferred similar diversity ratios and thus a similar degree of purifying selection for Meso European-like and Neo Aegean-like samples, albeit considerable individual variation. This suggests that the larger diversity seen among Neo Aegean-like samples is not simply the result of a larger effective population size, but rather of a particularly diverse source population, maybe as a result of past admixture (Marchi *et al*., 2022).

### Low X/A diversity of a Mesolithic female

To test for differences in sex-biased gene flow, we next compared the heterozygosity at neutral regions on the X chromosome and autosomes (X/A diversity) for all female individuals with whole-genome data (Fig 3D). Low X/A ratios may reflect demographic events such as recent bottlenecks or relatively low effective sizes for females compared to males (Pool and Nielsen, 2007; Amster and Sella, 2020; Amster *et al*., 2020). The lowest diversity ratio inferred was for the Early Mesolithic individual LEPE51, the only Meso European-like female for which sufficient sequence data is currently available. Among modern samples, equally low ratios were found for two individuals from forager populations (KhomaniSan-1, Saami-1). For other modern and all Neo Agean-like individuals we inferred higher ratios, albeit with substantial variation.

### Neolithic individuals (in the Danube Gorges) were smaller and tended to have lighter pigmentation than indigenous foragers

The genetic differences between local Mesolithic and incoming Neolithic populations at the Danube Gorges translated into observable phenotypic differences. Here, we focused on the four pigmentation phenotypes, *skin pigmentation*, *eye pigmentation, hair pigmentation* and *hair shade,* that we predicted using the HIrisPlex-S system. Focusing on non-admixed Meso European-like and Neo Aegean-like individuals along the Danubian corridor, none of these phenotypes were significantly different between these groups. In combination, however, these four pigmentation phenotypes were predictive of ancestry: using a linear discriminant analysis (LDA), we identified a combination of lighter skin pigmentation, bluer eyes, lighter hair pigmentation and lighter hair shade that predicted Neo Agegean-like ancestry for 30 (63%) of all 48 Neo Aegean-like individuals and two (40%) of the five admixed individuals with posterior probability > 0.75, while all 15 Meso European-like had posterior probabilities < 0.5 and nine (60%) even < 0.25 (Figure 4). The remaining 18 (37%) Neo Aegean-like individuals, however, appear to have had phenotypic combinations rather similar to Meso European-like individuals (i.e. darker pigmentation), as did the two second-generation admixed individuals from Lepenski Vir. Notably, Neo Aegean-like individuals from the Marmara region and Greece south of the Danube Gorges (the sites of Aktopraklık, Barcın, Nea Nikomedeia, Kleitos and Paliambela) were more easily distinguished from Meso European-like individuals (13/15 or 87%) than Neo Aegean-like individuals from the Danube Gorges or further north (17/33 or 52%, χ^2^=4.04, p=0.044). Nonetheless, more than half of all Neolithic immigrants could easily be told apart from local Mesolithic individuals just based on these four phenotypes, and likely even more based on the entire habitus.

**Figure 4.**
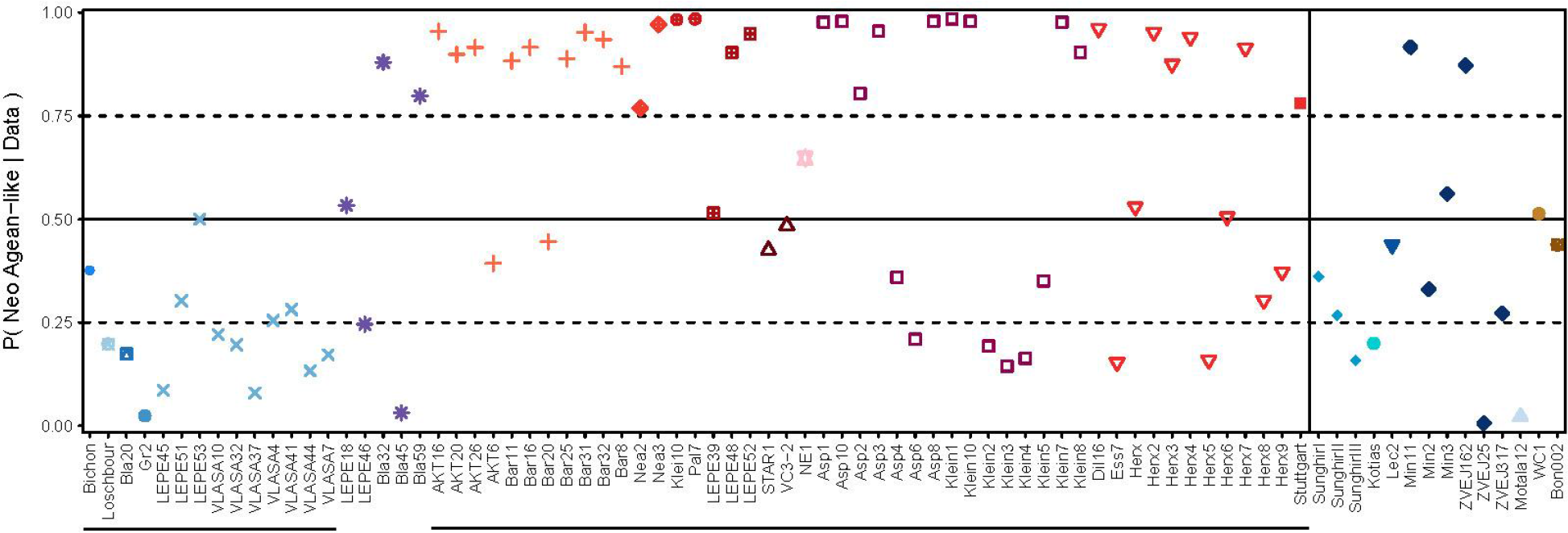
Posterior probabilities of HIrisPlex data based on a Linear Discriminant model (LDA). The model was trained on a set of samples (indicated by lines below x-axis) representing the Meso European-like and Neo Aegean-like populations, where the first discriminant function is a linear combination of four different phenotypes: skin pigmentation, hair pigmentation, eye pigmentation and hair shade. To reduce noise, we turned each phenotype into a binary configuration (see Methods). The trained model was then used to obtain posterior assignment probabilities for all samples, including those not used for training (the admixed samples and those from a different geographical context shown on the right).

A particular phenotype previously shown to differ between these groups, for instance, was a larger body size for Mesolithic than Neolithic individuals (Olalde *et al*., 2014; Ju and Mathieson, 2021; Marchi *et al*., 2022). Given the low number of Danube Gorges samples with reliable osteological estimates, evidence for differences in body size mostly stems from comparisons between periods, with Neolithic samples generally inferred as smaller than those from the Mesolithic period (Macintosh, Pinhasi and Stock, 2016; Jovanović, 2017; de Becdelièvre *et al*., 2020). However, it is interesting to note that the anthropological sexing errors discovered through our genetic analysis almost exclusively involved relatively tall Meso European-like females interpreted as males and relatively short Neo Aegean-like males as females (Supplementary Data Table 1, (Borić and Price, 2013; Budd *et al*., 2013; Roodenberg, Gerritsen and Özbal, 2013); personal communication).

## Discussion

### Genetic diversity in the Danube Gorges in a supra-regional context

On a projection-free PCA, Meso European-like individuals from the Danube Gorges spanning more than 1,500 years are well differentiated from other Meso European-like individuals, including those from Central Europe, North-Eastern Europe and the Black Sea region. This differentiation was initially hypothesized to be the result of admixture with more eastern hunter-gatherers (Mathieson *et al*., 2018; Feldman *et al*., 2019). However, it can also be explained by bidirectional gene-flow between different hunter-gatherer groups (Feldman *et al*., 2019), a hypothesis more in line with the deep split recently inferred between multiple hunter-gatherer individuals, including Meso European-like individuals from the Danube Gorges (Marchi *et al*., 2022). According to this scenario, the foraging communities of Europe suffered from a major population reduction and diverged into several smaller groups during the LGM. Members of one of those groups settled in the Danube Gorges, where they developed a semi-sedentary lifestyle certainly as of the Late Mesolithic period (Dimitrijević, Živaljević and Stefanović, 2016; de Becdelièvre *et al*., 2021). The level of gene flow with neighboring groups remains unknown. But the rather high pairwise diversity observed among the Danube Gorges individuals compared to, for instance, individuals from Central Europe, as well as the elevated heterozygosity of at least some Danube Gorges individuals is indicative of a relatively large and well connected population, an interpretation well in line with the richness of archaeological finds from this period (Borić and Stefanović, 2004; Borić *et al*., 2014; Borić, 2016, 2021). The difference in diversity appears particularly stark when compared to the genomic data from Criewen (GR2), the most recently dated Central European individual with 100% Meso-European-like ancestry, or the Baltic site of Zvejnieki. In contrast to the site of Zvienjeki, however, we found no indication of an influx of Mesolithic individuals with very distinct ancestries at the Danube Gorges.

That changed towards the end of the 7th millennium BC when individuals with Neo Aegean-like ancestry appeared in the Danube Gorges. The Mesolithic and Neolithic sites in the Danube Gorges provide a unique opportunity to genetically study forager-farming interactions at high resolution and interpret them in a European context. Whether Lepenski Vir was a forager community attracting individuals from the farming frontier, or whether the site was possibly newly occupied around c. 6,200 BC by arriving migrants from the Aegean region attracting forager individuals, has been a long-standing archeological debate (Srejović, 1969; Garašanin and Radovanović, 2001; Borić and Price, 2013). Signals of admixture between individuals with Meso European-like and Neo Aegean-like ancestry have been reported previously for several other prehistoric sites, where the rate of forager introgression was found to have been very low initially but to have increased significantly over time (Lipson *et al*., 2017). In contrast, evidence for farming individuals joining previously established forager societies remains rare (Bramanti *et al*., 2009; Bollongino *et al*., 2013; Hofmanová, 2016; Hofmanová *et al*., 2016; Mathieson *et al*., 2018), but early farmers, who were adapting their subsistence to new environmental conditions, might have been attracted by forager communities that provided local knowledge and access to resources, such as the permanent abundance in fish species in the Gorges at the time of the 8.2 cal BC cooling event. For local foragers, in turn, the resulting mutualistic relationships allowed for the exchange of goods and an access to a wider mating network (Bocquet-Appel and Bar-Yosef, 2008).

### Admixed individuals were also admixed culturally

This study brings evidence that admixture between individuals with Meso European-like and Neo Aegean-like ancestries certainly occurred before 6000 cal BC in the Central Balkans: the second-generation admixed individuals (male LEPE18, LV 27d; female LEPE46, LV 93), as well as the previously reported admixed case from Lepenski Vir (LEPI_61, LV 61; (Mathieson *et al*., 2018) were all dated to the period of Transformation (ca 6200-5950 cal BC). Although they were buried in extended supine position, following local foragers’ funerary customs, various contextual and bioarchaeological evidence indicate that admixture may have locally triggered a complex pattern of cultural mixing. For instance, the admixed male child LV 61 (6225-5915 cal BC; 4-7 years old) has been buried through the plastered floor of one of the oldest building from Lepenski Vir (building 40), in a practice with Neolithic Southern Balkans and Anatolian similarities (Borić and Stefanović, 2004; Stefanović and Borić, 2008). Mostly reserved at Lepenski Vir to infants and young children, and to a few adults with non-local strontium signatures (Borić and Price, 2013), this practice may have been associated with their social construction or status. According to stable isotope values, this child has been fed with large amounts of aquatic resources, consistent with nutritional socialization (de Becdelièvre, 2020). Similarly, the second-generation admixed female with ¾ Neo Aegean-like ancestry (LEPE46, LV 93; 6226-6026 cal BC) was also buried into a building (building 72) with various cultural elements pointing to the Early Neolithic cultural sphere (including numerous limestone beads, as well as a fragmented stone ring and adze). In contrast, the second-generation admixed male individual with ¾ Meso European-like ancestry (LEPE18, LV 27d; 6226-6026 cal BC) was discovered in a primary disturbed burial that contained grave goods associated with both Mesolithic (deer antler) and Neolithic (pottery fragments) communities. The other individuals, buried in extended or slightly flexed positions in continuity with local Mesolithic traditions, included the Meso European-like LEPE53 (LV 27a) and LEPE17 (LV 27b) with a mtDNA haplogroup frequently observed in early Neolithic farmers in Europe (N1a; Supplementary Data Table 1). All had a typically local diet, rich in aquatic proteins (Supp Info). The funerary practices associated with these individuals thus reflect the mosaic pattern of the Mesolithic and Neolithic cultural assimilation at Lepenski Vir.

### First-generation immigrants were culturally inter-connected

Some individuals, namely LEPE39 (LV 82), LEPE48 (LV 122) and LEPE52 (LV 73), likely represent first-generation immigrants that did not admix: they all had >96% Neo Aegean-like ancestry, had a diet distinct from that of local Meso European-like samples (Supplementary Data Table 1), and the two samples for which strontium was measured (LEPE52, LEPE48) showed non-local signatures (Borić and Price, 2013). Of those, the male LEPE52 from the Early Neolithic Period was buried in a flexed position, which is considered typical for Anatolian and Balkan Early Neolithic communities. However, the burial practices associated with the two samples from the earlier Transformation Period reflect elements likely associated with both Neolithic and Mesolithic funerary rites: The male LEPE39 has been discovered disarticulated and only the isolated skull (calvaria) of the non-local young (15-20 years) female LEPE48 was found beneath a building floor (building 47). The practice of disarticulating the skull, while also found among Early Neolithic Anatolian communities and, albeit more scarcely, in the Southern Balkans (Talalay, 2004), was common during the Mesolithic at the Danube Gorges: among the Meso European-like samples from the Danube Gorges studied here, four samples (VLASA4, VLASA10, VLASA32, VLASA41), all predating the appearance of Neo Aegean-like ancestry, were buried with signs of disarticulation. Being buried in a dwelling, on the other side, is a funerary practice frequently found in Early Neolithic Anatolian contexts (Adams and King, 2011; Brami, 2017). Together, these samples thus again attest to the gradual pattern of social integration between the groups as well as the cultural transformation triggered by this interaction at the Danube Gorges.

#### Interactions between foragers and farmers in the Danube Gorges

While the results shown above attest to a certain degree of cultural syncretism during the Transformation Period, this does not seem to apply to all sites in the Danube Gorges. All Danube Gorges individuals for which we estimate at least some Neo Aegean-like ancestry were buried at Lepenski Vir. A single additional individual with some degree of Neo Aegean-like ancestry was previously reported from Padina (PADN_4; (Mathieson *et al*., 2018)), the only other Danube Gorges site at which trapezoidal houses were found. In contrast, at the nearby site of Vlasac, the genome-wide ancestry of all seven individuals analyzed, as well as mtDNA haplogroups identified for additional 9 individuals, are all consistent with Meso European-like ancestry (Supplementary Data Table 1), in line with previous reports for Vlasac and Hajducka Vodenica (Mathieson *et al*., 2018). The majority of these samples likely pre-date the arrival of the Neolithic. However, at least seven individuals are confidently dated to the Transformation period, making it unlikely to have missed Neo Aegean-like ancestry if it was common during this period at those sites.

A few hundred years later, in the immediate vicinity of the Danube Gorges, the sites of Vinča-Belo Brdo and Starčevo were established (Whittle *et al*., 2002; Tasić *et al*., 2015; Porčić *et al*., 2021). The sites, both associated with the Neolithic Starčevo-Körös-Criş cultural complex and a subsistence more oriented towards the consumption of C3 plants (such as crops), meat/dairy products of domesticates and wild game (Filipović and Obradović, 2013; Jovanović *et al*., 2019; Stojanovski *et al*., 2020), paint a highly contrasting picture: For the only individual with whole-genome data available from these sites (Star1; (Marchi *et al*., 2022)), we estimate >96% Neo Agean-like ancestry, a signal confirmed by seven new mtDNA lineages (Supplementary Data Table 1).

Collectively, this observation suggests the continued co-existence (6200-5950 cal BC) of foraging and early farming communities if not at the same site, then at least in the same settlement area. Only in Lepenski Vir and possibly in Padina does the interaction take place at the same site -perhaps even into the Neolithic period. However, the Vlasac site was possibly no longer used as a settlement during the Transformation period, but only as an ancestral burial site by people with Meso European-like ancestry (Borić *et al*., 2014).

Mesolithic cultural elements disappeared gradually. Several elements of the Neolithic culture, such as domesticated animals, crop consumption and a typical Neolithic symbolic repertoire, appear at Lepenski Vir only after the Transformation phase when trapezoidal houses were also abandoned and the flexed position became the new dominant mortuary canon (Porčić, Blagojević and Stefanović, 2016; Blagojević *et al*., 2017; Jovanović *et al*., 2019; de Becdelièvre *et al*., 2020). This cultural change coincides with a significant population increase at Early Neolithic sites in the Central Balkans and is associated with a general population increase and a higher percentage of individuals with non-local isotope signals, suggesting a second wave of Neolithic immigrantion (Borić and Price, 2013; Porčić, Blagojević and Stefanović, 2016; Blagojević *et al*., 2017; de Becdelièvre *et al*., 2021). In line with this view, the three first-generation immigrants at Lepenski Vir (LEPE39, LEPE48 and LEPE52) were all found to be unrelated and most likely date to different generations (Table 1). In addition, we found a lower fraction of the typical Meso European-like mtDNA haplogroup (U5) among individuals dating to the Early Neolithic Period than among individuals dating to the Transformation Period (3/15 vs. 12/23). Hence, the Neolithisation of the Danube Gorges should not be understood as a straightforward process of acculturation or a sudden behavioral shift. Results rather reflect a mosaic picture of complex behavioral interactions and increased immigrations which triggered gradual socio-cultural changes within the framework of local economic and ecological continuity.

### Conclusion

The analyses presented here consolidate the picture of the Neolithisation of South-Eastern and Central Europe within the framework of a demic diffusion. Using heterozygosity estimates, we show the decrease in genetic diversity from sites in NW Anatolia to those in Central Europe resulting from the demic expansion along the archaeologically attested expansion route. At the genetic level, the interaction of early Aegean farmers with European hunter-gatherer groups along the expansion route has been demonstrated mostly indirectly: while most studies agree that about 2-6% of the genome of early Neolithic European people derives from admixture with hunter-gatherers during the Early Neolithic period, direct genetic evidence for hunter-gatherers in an early Neolithic context is limited to a single individual reported from the Körös site Tiszaszolos-Domaháza in Hungary (Gamba *et al*., 2014), an agricultural settlement at the frontier of the Neolithic Expansion that persisted for a few generations only. Considering the large number of individuals studied from Early Neolithic sites so far, the scarcity of individuals with predominantly Meso European-like ancestry and the complete absence of early-generation admixed individuals is remarkable. In this study, for instance, we newly analyzed 21 individuals with genomic data from typical Early Neolithic sites across Europe (Herxheim, Kleinhadersdorf, Dillingen-Steinheim, Asparn-Schlelz), but have not found a single individual that shows substantial Meso European-like ancestry. Cultural practices such as differentiated burial rites may be responsible for this. Another, equally plausible explanation would be that intermarriage was not tolerated at typical Neolithic core sites itself, but perhaps only in the periphery.

So is Lepenski Vir a model for an experimental outpost on the Neolithic expansion front? Could intercultural practices have been tried out here that Neolithic societies, with their “colonist ethos” and entrenched narrow cultural practices, did not tolerate on their own land (Lüning, 2000; Özdoğan, 2011; Shennan, 2018)? Even if agriculture was not possible in the Danube Gorges, a connection to agricultural communities must have existed, as the temporal distribution of nitrogen isotope ratios shows. Whether this connection was also accompanied by gene flow from Lepenski Vir to the Neolithic communities has not been shown yet, but seems likely. Thus, sites like Lepenski Vir could well have been extramural contact zones between hunter-gatherers and early farmers, and thus responsible for the introgression of hunter-gatherer ancestry into Neolithic communities. This would not completely invalidate the alternative “dead-end theory” according to which Lepenski Vir was merely a failed early Neolithic experiment with a modified way of life, for both may be true.

## STAR Methods

### Key Resources Table

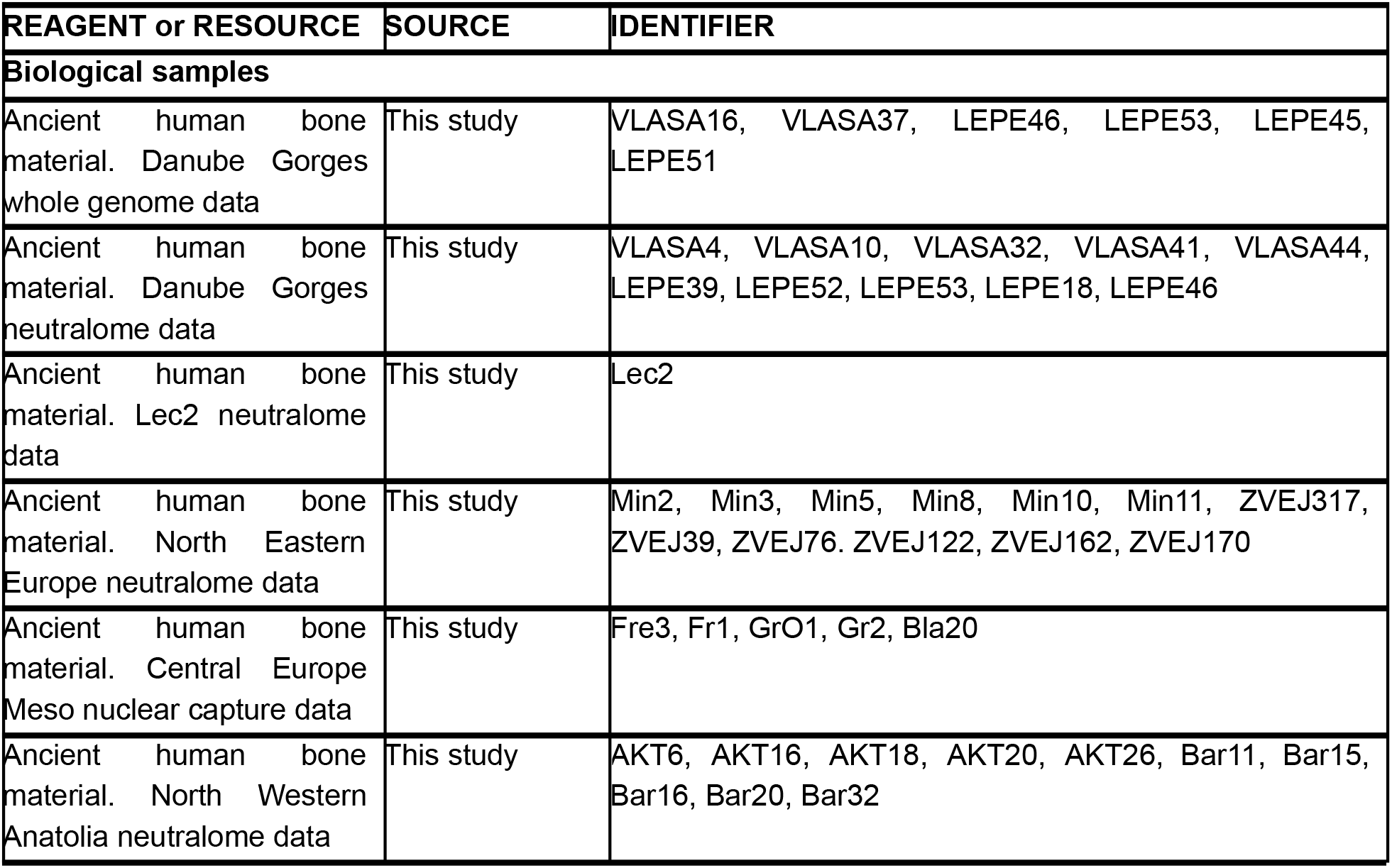
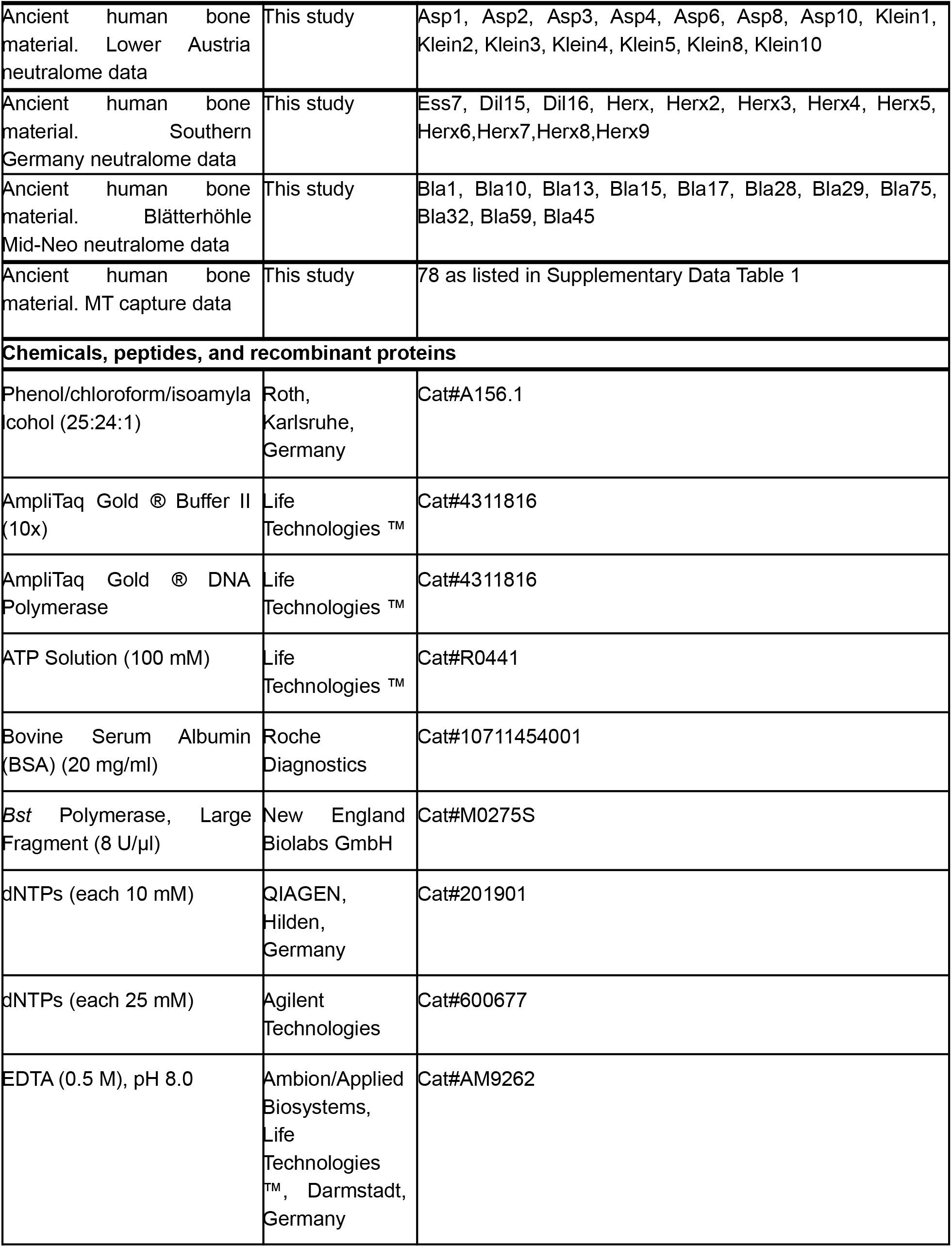
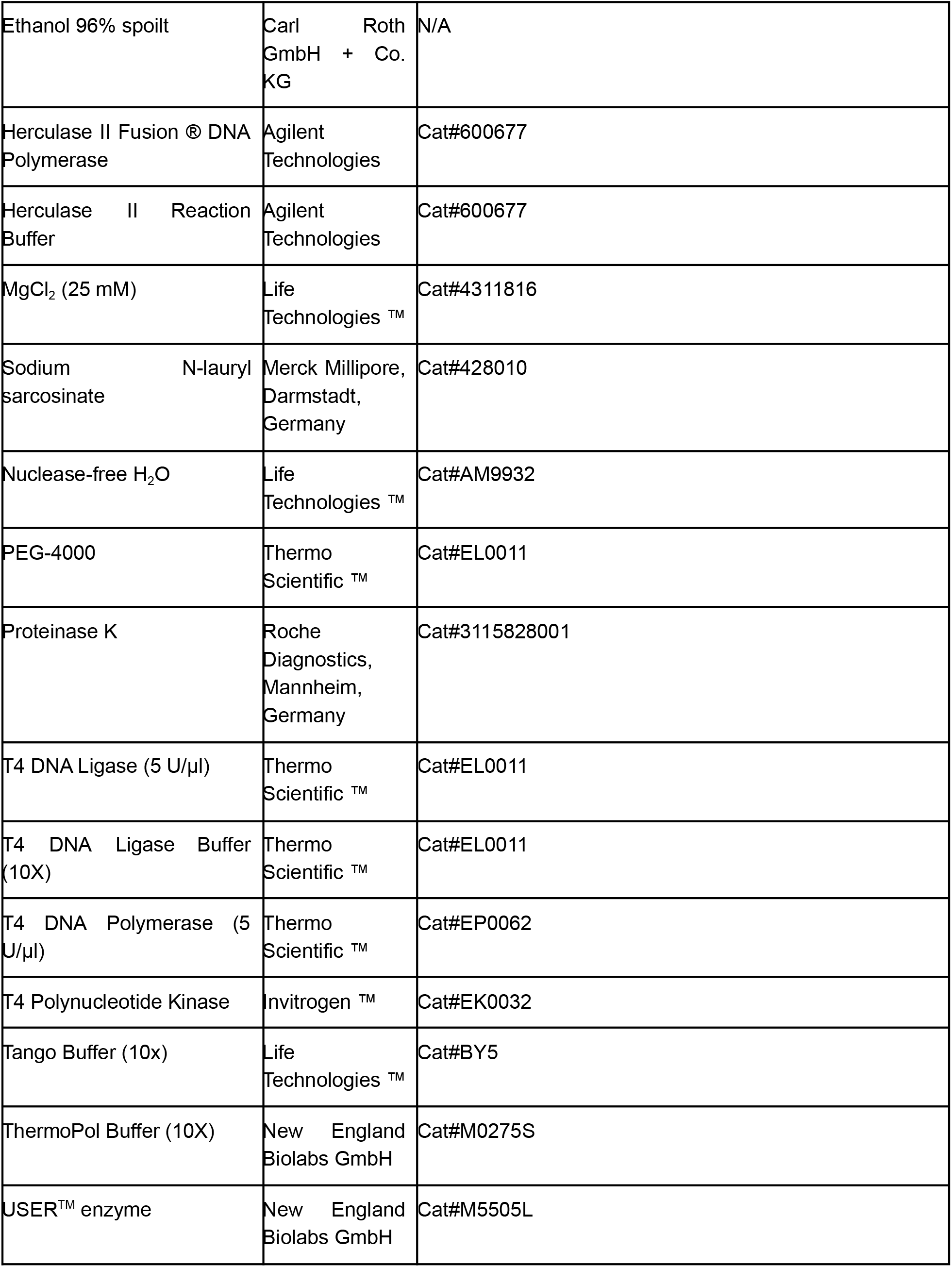
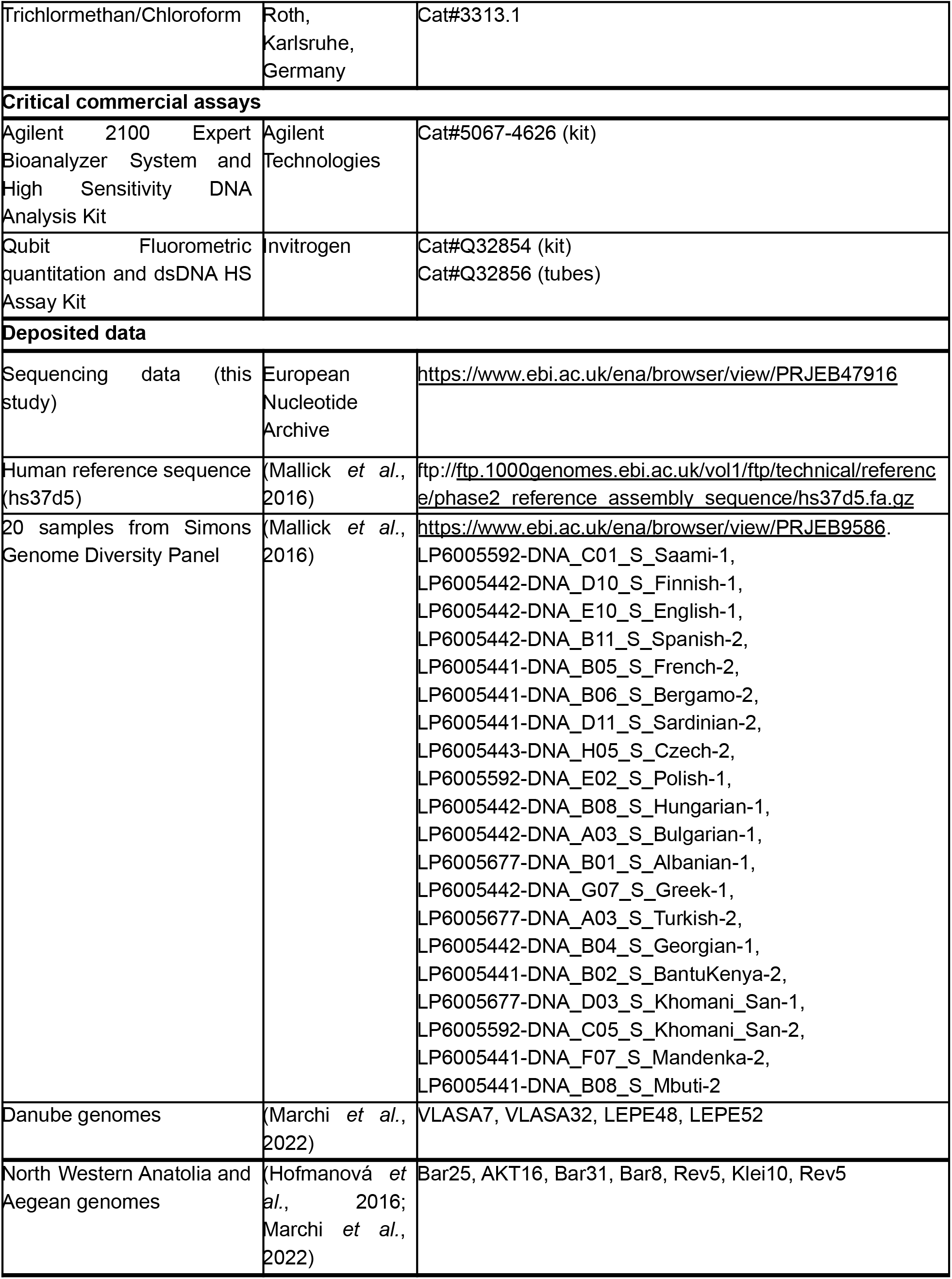
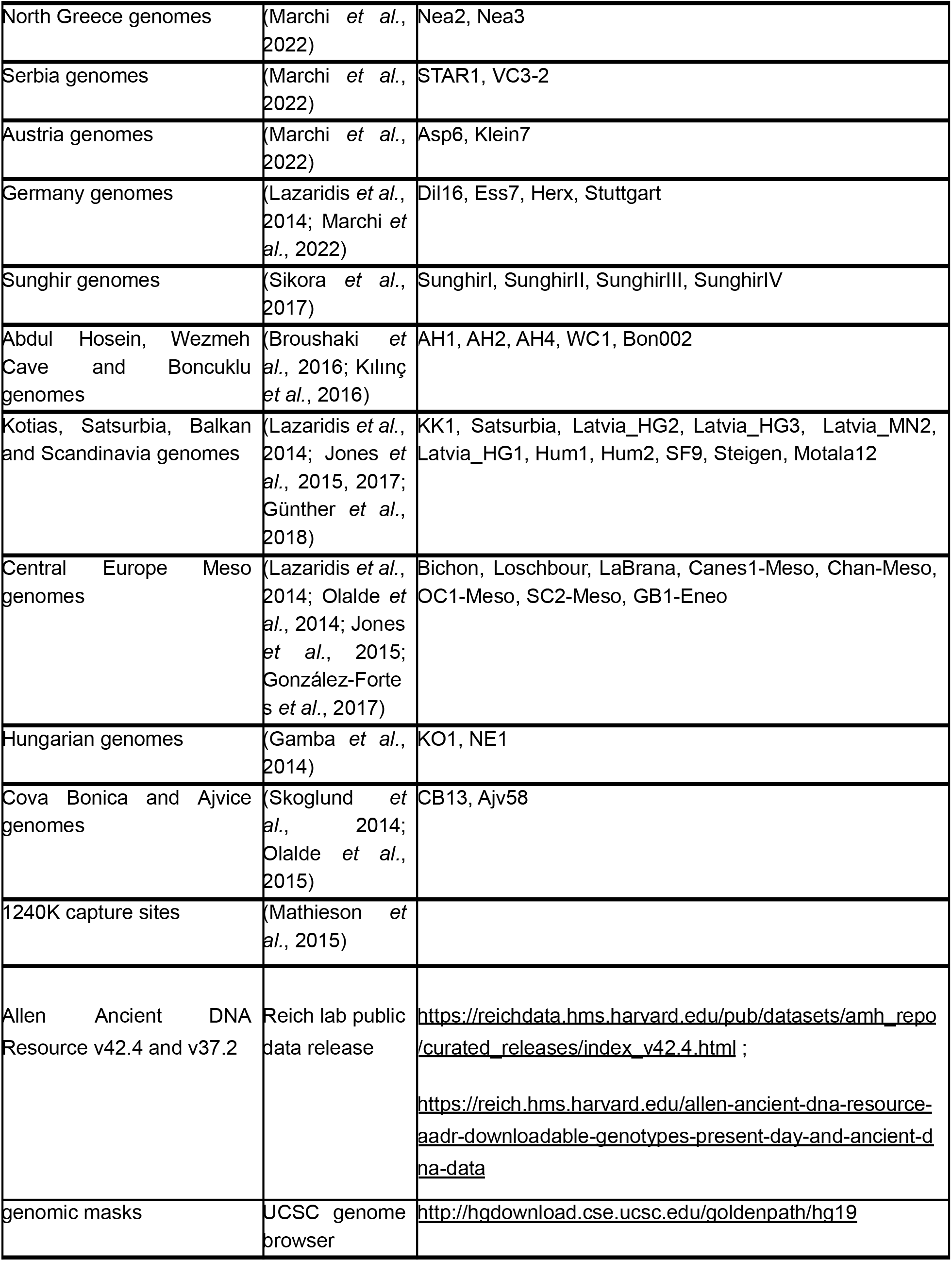
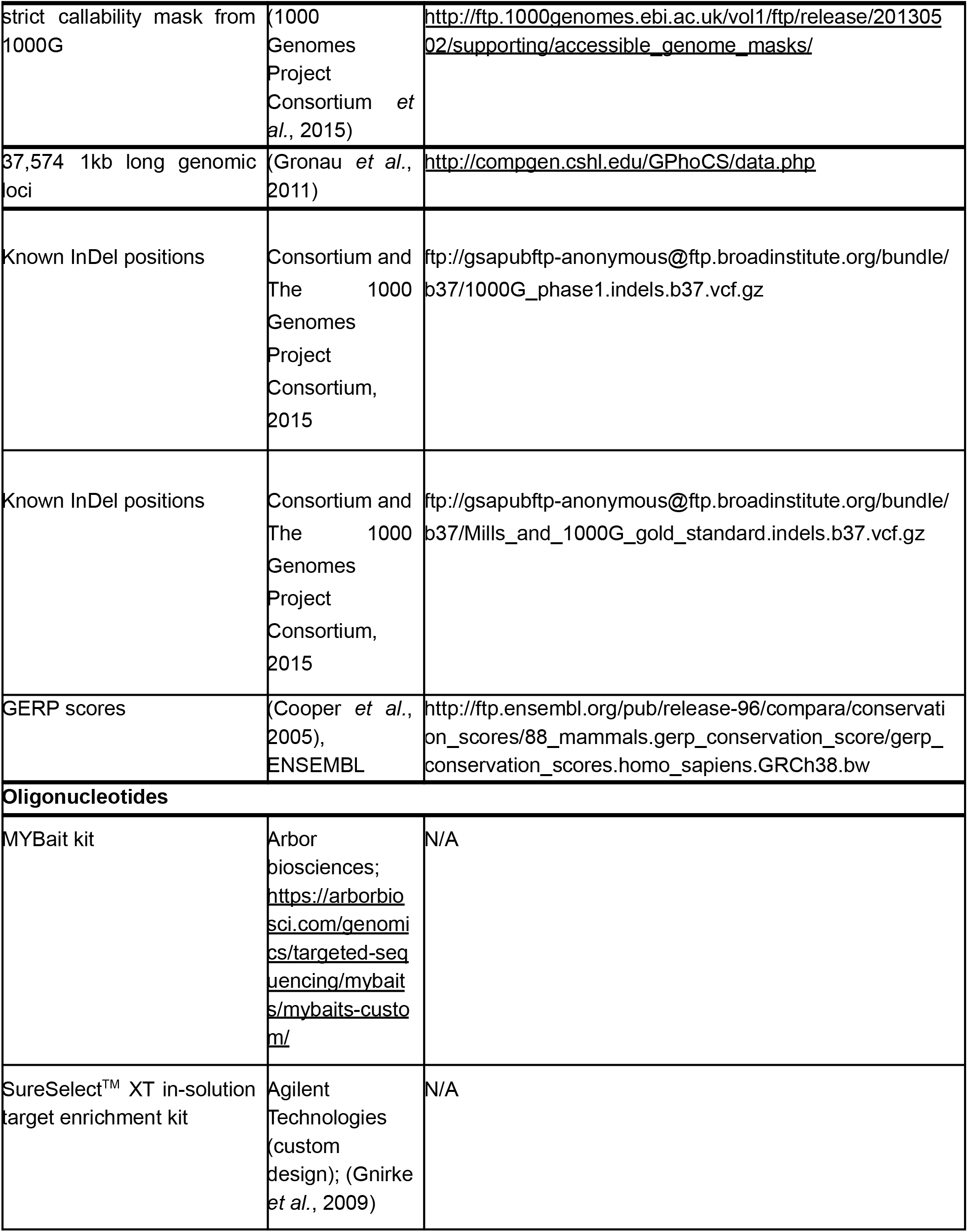
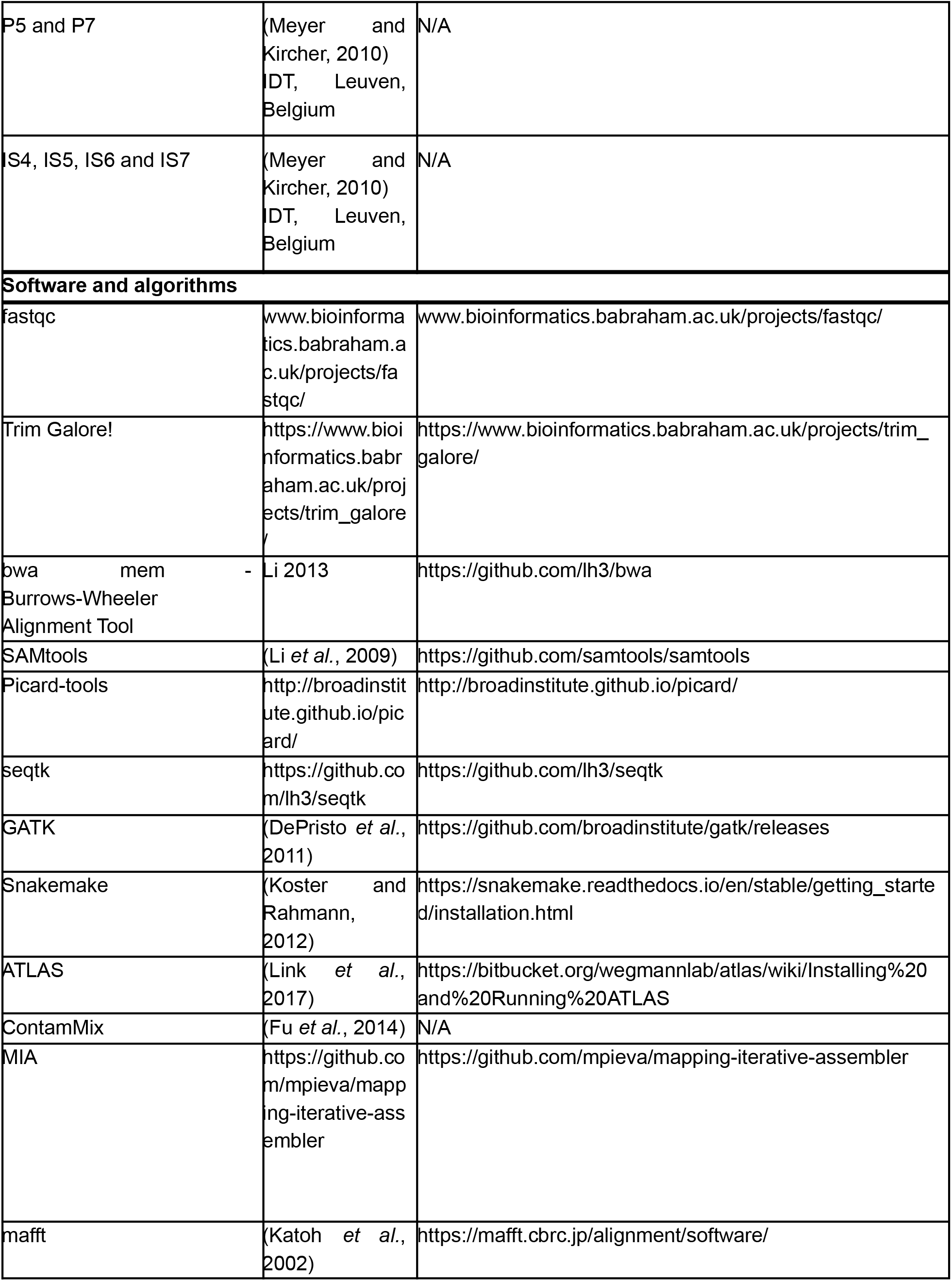
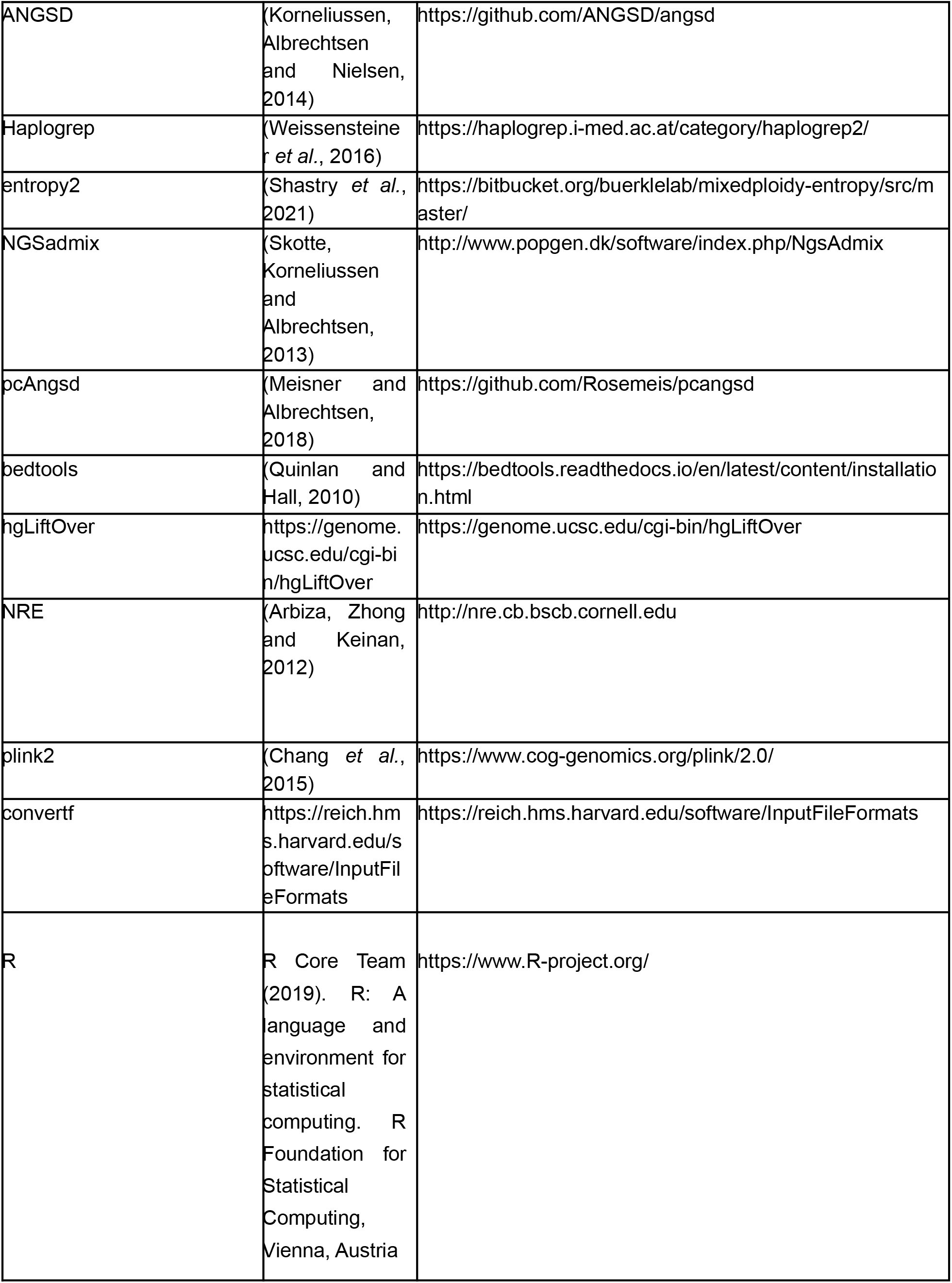
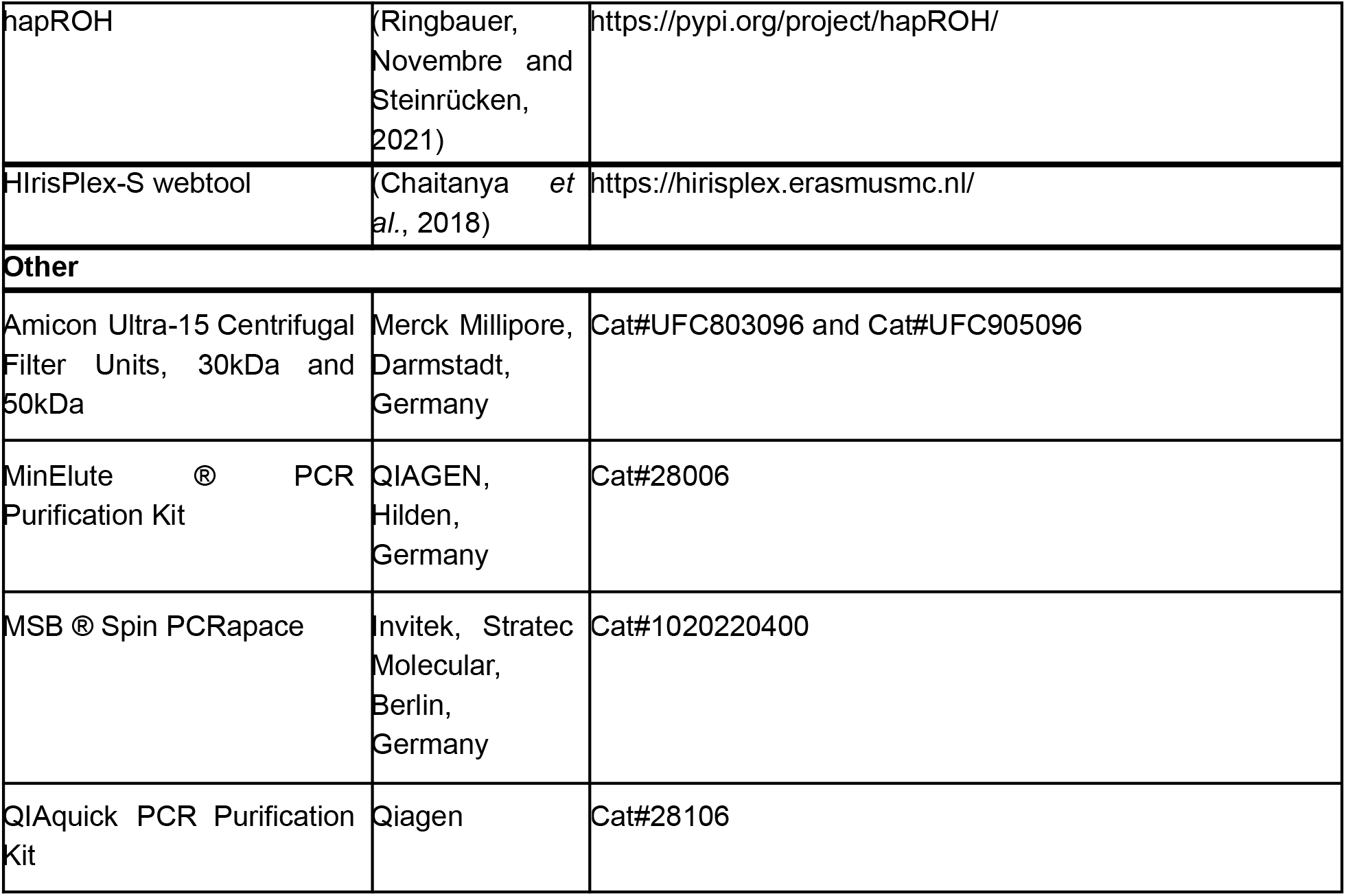

### Resource Availability

#### Lead Contact

Further information and requests for resources and reagents should be directed to and will be fulfilled by the Lead Contacts, Joachim Burger and Daniel Wegmann.

#### Materials Availability

Genomic data are available at the European Nucleotide Archive under the accession number PRJEB47916 in BAM and FASTQ format. Mitochondrial capture data are available at GenBank.

#### Data and Code Availability

All genomic data used in this study is publicly available at the European Nucleotide Archive under the accession number PRJEB47916 or at the sources listed in the Key Resources Table. The pipeline used to process the raw data is available on https://bitbucket.org/wegmannlab/atlas-pipeline/src/master/. The code used to plot relatedness is available on Bitbucket at https://bitbucket.org/wegmannlab/atlas/downloads/Relatedness.R.

### Experimental Model and Subject Details

#### Archaeological context

The samples analyzed in this study have been obtained from archaeological collections with permission of the responsible curators or excavators.

##### Barcın

The Barcın Höyük was occupied without interruptions between 6,600-6,000 cal BC (Gerritsen, Özbal and Thissen, 2013) and shows continuity from pre-Fikirtepe to Fikirtepe horizons of the Neolithic period ((Hofmanová *et al*., 2016); Suppl. S4-S5). While the site shares common elements with other Fikirtepe sites in the Marmara region, there were differences in architecture and dietary habits noted between flat sites (e.g., Aktopraklık) and tell sites (e.g., Barcın) (Karul and Avcı, 2013). This site is the oldest known Neolithic occupation in NW Anatolia (Gerritsen, Özbal and Thissen, 2013) and from the start, the food economy was fully agrarian with absence of an earlier transitional phase from foraging to farming (Arbuckle *et al*., 2014). The dead were buried at several locations within the settlement: neonates and infants were buried within or close to the houses, generally next to the walls, whereas juveniles and adults were buried in primary single burials in the central courtyard (Alpaslan Roodenberg, Gerritsen and Özbal, 2013).

##### Aktopraklık

Aktopraklık is a flat inland Fikirtepe site about 25 km from the city of Bursa (Marmara region) and its excavation showed uninterrupted occupation from the middle of the 7th millennium BC to the middle of the 6th millennium BC (Karul and Avcı, 2013). Site Aktopraklık C served as a settlement (with associated human remains) during the earlier phases and as a cemetery during the later phases (the settlement moved to Aktopraklık B during the Chalcolithic) (Karul and Avcı, 2011). In the Neolithic phase, the individuals were buried close to buildings or beneath them (Songül Alpaslan-Roodenberg and Roodenberg, 2020).

##### Asparn-Schletz

Asparn-Schletz is a large settlement enclosed by two ditch systems (Windl, 2009) with signs of occupation (Windl, 1999). There were 20 burials in the ditch and 130 individuals interred without a classical burial, often with perimortem trauma and likely buried some time after death (Teschler-Nicola, 2012). The burials predate the presumably violent event that has taken place in the end of the LBK, ca. 7150-6900 BP (Wild *et al*., 2004) and possibly the site was abandoned after this (Windl, 2009).

##### Kleinhadersdorf

The classical LBK cemetery, the largest such site in Austria, was located close to a LBK settlement and is dated to the second half of the 6th millennium BC (Neugebauer-Maresch and Lenneis, 2015). Features of the lithic technology and red ochre have been interpreted in the past as possible signals of continuation of Mesolithic traditions (Mateiciucová, 2015). Most graves were single primary inhumations in a contracted position on the left side typical of LBK with some exceptions (burials on the back, possible cenotaphs) (Neugebauer-Maresch and Lenneis, 2015; Tiefenböck and Teschler-Nicola, 2015).

##### Essenbach-Ammerbreite

Essenbach-Ammerbreite was an LBK cemetery close to an LBK settlement (Brink-Kloke, 1990). The same individual as in (Marchi *et al*., 2022) has been analysed in this study, namely Ess7, a child from grave 7.

##### Dillingen-Steinheim

Dillingen-Steinheim is a LBK cemetery of 27 classical burials (Nieszery, 1995), some in a ditch (Marchi *et al*., 2022). Two such individuals, analyzed in this study were Dil15 and Dil16 (graves 23 and 24, respectively) and they have been dated to 5,116 ± 118 cal. BC (Pechtl, 2015). They have been found to be brothers (see Results).

##### Herxheim

Herxheim is a well-known site, especially in relation to violence during the Early Neolithic period. The site is enclosed by two ditch segments that contained remains of more than 500 individuals (Haack, 2016; Zeeb-Lanz, 2016, 2019) with signs of perimortem and postmortem violence on the skeletons (Boulestin and Coupey, 2015). Interestingly, individuals formally buried showed a local range of Sr values in contrast to the scattered remains that showed more often a non-local signal (Turck, 2019).

##### Lesnik cave

The site of Lesnik cave is located near Yalta, Ukraine. The analyzed sample (Lec2) was not directly dated, but based on the contextual finds, Lec2 was placed in the early Mesolithic or Late Palaeolithic of the region. The other human sample (Lec1) from this cave, was dated to 11260+/-45 BP (OxA-19112), cal Oxcal 4.4 11292-11144 BC (Ridush, 2009; Schulz, 2016).

##### Zvejnieki

Zvejnieki in northern Latvia is a cemetery where individuals were buried from the middle Mesolithic to Neolithic period, from the 8th to 4th millenia BC (Eriksson, Lõugas and Zagorska, 2003; Stutz, Larsson and Zagorska, 2013). Individuals at the site were partially mobile and the inhumations did not respect one another, suggesting the absence of knowledge of previous burial placement in some later periods (Larsson *et al*., 2017).

##### Minino

While the site of Minino, Russia, is dated from the Palaeolithic to Neolithic, the main part is assigned to the Mesolithic period when burial complexes are very rare. The ages of the samples range from 5650-4600 cal BC to 8671±48 cal BC (Wood *et al*., 2013).

##### Criewen

Criewen, discovered in northern Germany, provided this study with samples associated with a non-agricultural context that were dated to 4770±40 cal BC (Gr1) and 4600±60 cal BC (Gr2) (Geisler and Wetzel, 1999). One of the samples was buried with ca. 3,000 perforated shells (Street *et al*., 2001).

##### Groß Fredenwalde

At Groß Fredenwalde, at least eight burials have been found with 12 individuals. The site was dated from 6,400 cal BC to 4,900 cal BC with one individual overlapping with the presence of farmers in the region in approx. 5,200 cal BC (Terberger *et al*., 2015; Kotula, Piezonka and Terberger, 2020).

##### Große Ofnet Höhle

33 skulls separated from their postcranial skeletons have been found at the site. They date to 5770 cal BC (Walde *et al*., 1986; Orschiedt, 1998).

##### Blätterhöhle

Blätterhöhle shows signs of human occupation from the Mesolithic to Late Neolithic. During the Mesolithic, the site has been used as a sporadic settlement. Human remains have been discovered inside the cave in a disturbed context, showing presence of hunter-gatherers and farmers through time but also in parallel (human remains from individuals with different subsistence were differentiated by isotopic analysis) (Orschiedt *et al*., 2012; Bollongino *et al*., 2013).

##### Lepenski Vir

This large and famous settlement in the Danube Gorges is a type-site of the Lepenski Vir culture (sometimes called Lepenski Vir-Schela Cladovei culture). The site with its abundance of burials has been under discussion ever since its excavation (started 1965 by Dragoslav Srejenović) and the chronology of the area has been revised with the use of radiocarbon dating corrected for freshwater reservoir effect (Borić, 2002). The sampling strategy and further details about the site are provided in other parts of this study.

##### Vlasac

The Vlasac site is geographically very close to Lepenski Vir (cca 3 km downstream) and was first excavated in the same period as Lepenski Vir (Boroneanţ, 2011). The site was assigned to the Lepenski Vir culture and is mostly dated to Late Mesolithic, while there are dates as old as 9,800 cal BC known from the site (Bonsall *et al*., 2000; Borić and Stefanović, 2004; Borić, French and Dimitrijević, 2008). Additionally, new excavations (during seasons 2006–2009) showed that there was also an occupation parallel to the Transformation phase of Lepenski Vir with appearance of features influenced by the Neolithic (Early Starčevo ceramics, Spondylus shells and discoid beads) (Borić *et al*., 2014). Most of the settlement was abandoned ∼6,300/6,200 cal BC, but the site might have served as an “ancestral” place and a cemetery for burial rites of Mesolithic tradition during the last centuries of the 7th millennium BC (Borić *et al*., 2014).

##### Grivac

This well-stratified Neolithic site in central Serbia contains both proto-Starčevo and Vinča layers (Bogdanović, 2008; Porčić, Blagojević and Stefanović, 2016). The individual included in this study (Gri1) was buried in grave 1.

##### Padina

Padina is a Danube Gorges site with a superposition of Mesolithic and Early Neolithic structures. It is associated with the Lepenski Vir culture, with additional later layers of Late Neolithic cultures present at the site (Borić and Miracle, 2004). The only ancient individual successfully analysed from this site (Pad 11, burial 30) is assigned to the Early/Middle Mesolithic (∼9,500-7,400 cal BC) (Borić and Price, 2013).

##### Ostrovul Corbului

Ostrovul Corbului is located on the Romanian side of the Danube Gorges, downstream from Lepenski Vir. It was originally assigned to the Schela Cladovei culture, which was later connected to the Lepenski Vir culture (Boroneanţ, 2011). There are Mesolithic and Neolithic layers at the site (Roksandić, 1999).

##### Vinča-Belo Brdo

Vinča-Belo Brdo is a typical site of the eponymous Vinča culture with key Neolithic developments such as the formation of large settlements and tells, the intensification of farming subsistence and the expansion of material networks (Tasić *et al*., 2015). The culture occupied a large region of Serbia and several bordering countries between the late 6th millennium BC and middle 5th millennium BC (Borić, 2015). The only burial at the site is a collective burial at the lowermost level (Dimitrijević, 2014) and it is still under discussion whether this Starčevo level represents continuous occupation to later phases or an earlier abandoned settlement (Borić, 2009). The collective burial can be dated to 5,476-5,304 cal BC (Borić, 2009) and the samples analyzed here are coming from this context.

##### Sultana Malu Roşu

This Eneolithic site is located on the bank of the Mostiştea River, about 15 km from the Danube (approximately 500 km from the Danube Gorges) (Lazăr *et al*., 2008).

### Method Details

#### Sample preparation

Sample preparation took place in a dedicated ancient DNA facility of the Paleogenetics Group at the Institute of Organismic and Molecular Evolution (iomE) at the Johannes Gutenberg-University in Mainz, following and further improving guidelines for good practices in ancient DNA analysis (Bramanti *et al*., 2009; Bollongino *et al*., 2013; Scheu *et al*., 2015). We processed blank controls alongside milling, extraction and library built to control for the decontamination procedure of the devices used. Sample treatment and library preparation were performed as described in (Kircher, Sawyer and Meyer, 2012) with the adaptations described in (Scheu *et al*., 2015) and (Hofmanová *et al*., 2016). Sample specific modifications/adaptations are noted in Supplementary Data Table 1. 12 samples underwent a pre-lysis step during DNA extraction as described in (Scheu *et al*., 2015) in order to increase the percentage of endogenous yields. USER^TM^ (NEB) treatment of the DNA extract was performed for 181 libraries prior to Library preparation. To increase library complexity, we amplified each library in three or six PCR-parallels.

#### Quality assessment

With the first library of each sample, we estimated library complexity by quantitative real time PCR (qPCR) as described in (Meyer and Kircher, 2010; Hofmanová *et al*., 2016). Additionally, we performed a shallow screening on an Illumina Miseq (50 bp, single end sequencing) that was analyzed with the pipeline described in (Hofmanová *et al*., 2016).

#### Library preparation and Target enrichment

Based on the endogenous DNA content (fraction of reads aligning to the reference genome) and the qPCR results (Figure S1A) we chose three strategies on a per sample basis, for which additional extractions and libraries were created: 1) We chose samples of highest quality for whole genome sequencing, aiming to obtain genomes of differing ages from the Danube Gorges. Selected samples were sequenced on an Illumina Hiseq2500 (100bp paired-end or single-end) and an Illumina NextSeq (75bp paired-end). 2) We also selected a total of 75 high quality samples to perform a nuclear target enrichment (nuclear capture). The capture array of 5Mb sites (i.e. neutralomes) followed the protocol as described in (Veeramah *et al*., 2018) with some alterations. Sequencing was performed on an Illumina Hiseq 2500 with 100bp single-end or paired-end runs and on an Illumina NextSeq with 75bp paired-end sequencing at the University of Mainz. 3) We performed mitochondrial target enrichment for samples with relatively low DNA quality. Target enrichment was performed two times (double capture) following the methods described in (Hofmanová *et al*., 2016), with the exception, that PCR purification was performed with QIAquick PCR Purification Kit columns (QIAGEN^®^) according to the companies protocol, eluting in 33µl preheated (65°C) elution buffer for 5 minutes. For 9 samples (Supplementary Data Table 1), the supernatants from both nuclear target enrichment steps containing non-target DNA, were used to perform an additional mitochondrial capture experiment (supernatant capture). The supernatants were purified with QIAquick PCR Purification Kit columns prior to the first mitochondrial target enrichment. Samples were pooled equimolarly and sequenced at GENterprise GENOMICS (Mainz) on an Illumina MiSeq sequencing system with 50 cycles single-end or on an Illumina HiSeq sequencing system with 100 cycles paired-end. Mitochondrial data was additionally gained as by-products of the previous two strategies.

#### Bioinformatic processing

The mitochondrial captures were processed as described in (Hofmanová, 2016; Hofmanová *et al*., 2016). Read statistics can be found in Supplementary Data Table 1. For all 134 ancient genomes and neutralomes, alignment, local realignment, PMD and recal estimation were processed with commit *9ec713b* of the ATLAS-Pipeline (bitbucket.org/wegmannlab/atlas-pipeline/wiki/Home) with some minor changes depending on how the data were obtained as indicated below and in Supplementary Data Table 1. Reads were trimmed with length filter ≥ 30 (*TrimGalore*, v0.6.4, https://github.com/FelixKrueger/TrimGalore), aligned with *bwa-mem* (v0.7.17, (Li, 2013)) to the hs37d5 reference (Mallick *et al*., 2016), filtered for mapping quality < 30, sorted and indexed (*SAMtools*, v.1.9, (Li *et al*., 2009)). Read groups were added with *picard-tools* (v2.21.1, http://broadinstitute.github.io/picard/) to keep track of libraries. Unmapped reads, orphans and secondary alignments were removed with *SAMtools*, and duplicates were marked with *picard-tools* before and after merging the library BAMfiles per sample. For neutralomes, we merged all BAMfiles into one *master* BAMfile for further processing. A custom script was run to download data, mark duplicates and filter out reads as above for 20 SGDP BAMfiles (Mallick *et al*., 2016). All samples underwent Local Realignment (*GATK*, v.3.7; (DePristo *et al*., 2011)) using a union interval set of 30 samples plus the target sample (*RealignerTargetCreator*), and a guidance set of 12 samples for realigning along the target sample with *IndelRealigner* (Supplementary Data Table 2). We used *ATLAS* (v0.9*; (Link et al., 2017)*) with commit 7c1e6a4, unless indicated otherwise, and filterSoftClips option to split/merge single-end/paired-end reads (task=splitMerge) which generated our final processed BAMfiles. See Table 1 and S1 for more information.

### Quantification and Statistical Analysis

#### Library and Sample Statistics

We used *ATLAS* (task=BAMDiagnostics and task=depthPerSiteDist) and *SAMtools* flagstat to determine read counts, sequencing depth, endogenous DNA-content and further statistics that are listed in Supplementary Data Table 1 for each library parallel and merged samples.

#### Molecular Sex Determination

Using the script by (Skoglund *et al*., 2013) obtained from https://github.com/pontussk/ry_compute), we determined the molecular sex of individuals by calculating the ratio of reads aligned to the Y chromosome over the total number of reads aligned to X and Y. Ratios of 0.075 or higher indicate males and ratios below 0.016 indicate females (Figure S1C). It was run with default settings as suggested by Skoglund’s documentation.

#### Ancient DNA Authenticity

The blank controls from milling, extraction, library and capture experiments were measured by Qubit® Fluorometric quantitation (dsDNA HS assay, Invitrogen, Carlsbad, California, United States) and on an Agilent 2100 Bioanalyzer (HS DNA, Agilent Technologies, Waldbronn, Germany). Mitochondrial capture controls were sequenced alongside the capture sequencing. The concentration of potential contaminants was never higher than 2.1 ng/ul and the screening results showed a maximum of 39 aligning reads per blank control (out of a potential share of 80.000 reads). PMD-patterns were checked for all samples to show the characteristic exponential pattern of ancient DNA (Figure S1B). We estimated contamination on mitochondrial regions with *ContamMix (*v1.0; (Fu *et al*., 2014)) and for male individuals on the X-chromosomal regions with ANGSD (v0.917-108-g2f9cc4b; (Rasmussen *et al*., 2011; Korneliussen, Albrechtsen and Nielsen, 2014)). As the nuclear capture regions do not span the mitochondrial DNA and only contain few regions on the X-chromosome, we require a minimum of 10X depth over the mitochondrial genome (to assure correct consensus-calls) combined with a mt/nuc ratio below 200 (Furtwängler *et al*., 2018), as well as a minimum of 100 SNPs for ANGSD estimation to rely on the results of contamination estimation (Nägele *et al*., 2020). The detailed results can be found in Supplementary Data Table 1.

#### mt-DNA analysis

##### mt-Capture of nuclear capture supernatants

An additional mitochondrial target enrichment experiment on the supernatant of a nuclear target enrichment experiment - meaning the DNA not hybridized on the nuclear target - yielded a significantly higher percentage of endogenous reads aligning to the mitochondrial genome than the conservative mt-capture (t-test 4.0, p=2.3e-04). This could be explained by a higher percentage of mitochondrial fragments in the hybridisation process as well as a potential lack of steric hindrance in the hybridisation process as much longer nuclear molecules have been removed. Yet, the fold-coverage is significantly lower for supernatant captures (t-test −6.5, p=5.75e-08). This is expected, as several additional purification steps are performed on supernatant captures, accompanied by a severe loss of molecules. As a supernatant capture will only be performed after a nuclear capture, and therefore on high-quality samples, it can be recommended for further experiments to reduce the amount of sample material used.

##### mt-DNA Haplogroups

In order to determine the mitochondrial haplotypes from nuclear genomic data, majority allele calls on the MT genome were created with ATLAS (task=call method=majorityBase). The output VCF files were merged and uploaded to the HaploGrep 2.0 Website (Weissensteiner *et al*., 2016). The mitochondrial haplotypes for mitochondrial capture experiments were obtained as described in (Hofmanová, 2016; Hofmanová *et al*., 2016).

#### Genotype likelihoods estimations

##### Post mortem damage patterns

We used *ATLAS* (task=PMD) to infer position-specific PMD patterns as described in (Kousathanas *et al*., 2017) from the tabulated mismatches between the raw reads and the hs37d5 reference genome. These patterns were mainly inferred independently for all whole-genome individuals and read groups. However, for neutralomes we used the *master* BAMfile to pool all read groups that came from the same sample and had similar PMD pattern (option *poolReadGroups*).

##### Base quality score recalibration

We used *ATLAS* (task=recal) to recalibrate base quality scores with the method described in (Kousathanas *et al*., 2017) using the model *qualFuncPosSpecificContext*. This reference-free approach exploits a set of known homozygous sites and is extended to additional covariates beyond the original quality score, in our case the specific position within the sequencing read and the nucleotide context. As known homozygous sites, we used 10 million sites highly conserved among mammals as reflected by high RS-Scores (also called GERP scores; (Cooper *et al*., 2005)) calculated across the multiple sequence alignments of 88 mammals and provided by Ensembl (http://ftp.ensembl.org/pub/release-96/compara/conservation_scores/88_mammals.gerp_conser vation_score/gerp_conservation_scores.homo_sapiens.GRCh38.bw). For single-end sequencing data, the read groups were split into two new read groups during the raw data processing, so we provided the names of these split read groups to be merged for recalibration (option *poolReadGroups*). For neutralomes, we used the *master* BAMfile to pool all read groups that were generated with the same sequencing run and lane. This increased the power when estimating recalibration parameters for all neutralome samples.

##### Genotype likelihoods

We used *ATLAS* (task=GLF) to infer genotype likelihoods for all individuals at autosomal neutral capture 5Mb sites. We considered the PMD and the recalibration parameters previously estimated. This created GLF files for all our samples at the selected sites.

#### Downstream quality filtering

##### Downsampling experiments

For further analysis, we tested the impact of recalibration and post-mortem damage in 12 low-coverage and medium-coverage whole-genome samples. The tests consisted in downsampling the BAMfiles by probabilities ranging from 1 to 0.05, and estimating their heterozygosity in windows (*ATLAS*, task=thetaQC), either ignoring or considering PMD and recalibration parameters, respectively. For more details, see Figure S2B.

##### Depth and Heterozygosity Filtering

Based on the downsampling experiments, we estimated heterozygosity once with full data and 50 times using a downsampling probability of 0.5 (*ATLAS*, task=thetaQC) for every sample. The PMD and recalibration parameters inferred from full data were used in all heterozygosity estimations so that we can assess how well the error-rates are recalibrated between the full data and downsampled versions. We calculated the log ratio between the median sampled estimate with full data and the median of the 50 downsampling median sampled estimates. For the neutralome data, we bootstrapped 100 times the genome wide heterozygosity on the capture neutral regions and took the medians for the full data and the 50 downsampling median estimates and proceeded to calculate the log ratio. A ratio <0.239 and >-0.239 and a depth of coverage >1.5x for whole-genome and >4x for neutralome were used as quality filters (Figure S2A) to ensure comparability among samples and increase the sensitivity of our population genomic analyses. A total of 54 out of 79 whole-genomes and 56 out of 75 neutralomes passed these filters (7 overlaps between them) with two neutralome samples being further removed as mentioned below (Kinship analysis).

#### Population genetic analysis

##### Kinship analysis

We used *ATLAS* (task = geneticDist) to estimate the euclidean distances between all pairs of ancient samples that passed our quality control filters, using the GLFs created during the estimation of genotype likelihoods. We then used a custom R script (https://bitbucket.org/wegmannlab/atlas/downloads/Relatedness.R) to apply the method of (Waples, Albrechtsen and Moltke, 2019) to transform these distances into estimates of genetic relatedness. No relatedness was detected among Danube Gorges samples, but for Dil15 and Bar15 which were filtered out for downstream analysis. See details in Figure S1D.

##### Major/minor

We used *ATLAS* (task = majorMinor) to estimate the major and minor alleles (default parameters) from our sample-specific GLFs and output the genotype likelihoods for those in a vcf file, which then was converted to a Beagle format (task=VCFToBeagle) for the population genetic analysis. This was run for different sets of populations.

##### PCA

We used *PCAngsd* (v0.986; (Meisner and Albrechtsen, 2018)) with default parameters (MAF>=0.05) on different sets of populations to estimate the covariance matrix and perform Principal Component Analysis (PCA).

##### Admixture analysis

We used *NGSAdmix* (Skotte, Korneliussen and Albrechtsen, 2013) with -minMaf 0.05 to infer admixture proportions for different numbers of clusters (K=2 to K=7). Each K was run 10 times with a different seed and the best K was estimated by applying the Evanno method (Evanno, Regnaut and Goudet, 2005). In addition, Entropy (v2.0; (Shastry *et al*., 2021)) and PCAngsd were run as well for different Ks to estimate admixture proportions. The deviance information criterion (DIC) was used for entropy models with K=2 to K=7, lower values of DIC correspond to better model fit. For PCAngsd, a different eigenvector value was provided each time since the best K was empirically set to 2 based on PC loadings. The admixture proportions were then compared among the three (Figure S3B). The results were qualitatively the same, in all cases K=2 is the best fit, and only the proportions from *NGSadmix* were used for population structure.

##### Inter population ancestry

We used the second model of *Entropy* (Shastry *et al*., 2021) that takes into account the combination of ancestry states across all loci in diploid individuals. The benefit of this model is that it allows distinguishing among early generations of admixed individuals (i.e., F1, BC1) when combined with their global ancestry. We followed the user manual as described in the bitbucket repository (https://bitbucket.org/buerklelab/mixedploidy-entropy/src/master/vignette_entropy.pdf) to format the input genotype likelihood data and run *Entropy* with the right options. The model was run with five chains simultaneously, where posterior distributions were estimated with 100,000 iterations, sampled every 10th iteration and with 10,000 burn-in.

#### Intra-inter genetic diversity

##### Genomic regions of interest

We extracted exons and introns from the human genome annotation file (http://ftp.ensembl.org/pub/grch37/release-104/gff3/homo_sapiens/Homo_sapiens.GRCh37.87.gff3.gz) by using a custom R script. We used *bedtools* (v2.27.1; (Quinlan and Hall, 2010)) to merge intervals and to subtract the exons regions that were also in the introns. We combined tracks from build GRCH37 of the UCSC Genome Browser (Kent *et al*., 2002) to exclude simple-repeats, segmental duplications, self Chains, regions with high CpG content and the strict callability mask from 1000G (1000 Genomes Project Consortium *et al*., 2015). We ended up with ∼50Mb in both regions (introns were downsampled to the same amount of exons sites). Additionally, we identified 17,737 neutral 1kb autosomal loci based on 37,574 autosomal neutral regions (Gronau *et al*., 2011) that were lifted over from hg18 to GRCH37 (https://genome.ucsc.edu/cgi-bin/hgLiftOver). To update and provide an extra layer of stringency while accounting for the liftover, we used *NRE* (Arbiza, Zhong and Keinan, 2012) to mask all sites that do not fall in the 37,574 lifted loci and removed mammalian conserved noncoding elements plus 100bp each side (PhastConsElements46WayPlacental GRCH37 track; (Pollard *et al*., 2010)). Regions with recombination rate <0.1 and >10 cM/Mb, nearest gene distance <0.01 cM, simple repeats, BG selection coefficient <0.85 and not separated at least 50kb from each other were excluded too following (Gronau *et al*., 2011; Veeramah *et al*., 2018). Since the list of 37,574 loci does not include the X chromosome, we applied the filters from above and (Gronau *et al*., 2011) on the X chromosome with slight modifications to be less stringent. We ended up with 5,505 neutral >1kb X loci (∼11Mb).

##### Genome-wide Heterozygosity (θ)

We used *ATLAS* (task=theta thetaGenomeWide minDepth=2 bootstraps=100) to infer genome wide heterozygosity with 100 bootstraps on autosomal neutral sites. The nuclear capture data in fact contain 5Mb of autosomal neutral sites at high depth; however, for ancient whole-genomes, more neutral sites were necessary for increasing the sensitivity of the estimates. Hence, we provided the autosomal neutral ∼18Mb sites (section “Genomic regions of interest”) for the whole-genome data and the autosomal neutral 5Mb sites for the nuclear capture data to estimate single genome-wide estimates for θ.

##### Heterozygosity ratio (θ_1_/θ_2_)

We first tested the statistical power of the heterozygosity ratio model in *ATLAS* by simulating genomic data with default parameters (task=simulate) as shown in Figure S4. It was concluded that a depth >1.5x and a window size >10Mb for both regions of interest provide good estimates. We then used *ATLAS* (task=thetaRatio) to estimate genetic diversity between exons and introns with a prior=0 for all whole-genome samples, and also between X and autosome neutral sites with a prior=log(¾) for all female whole-genome samples. These genomic regions are above 10Mb and they were extracted as explained in section “Genomic regions of interest”.

##### ROHs

We used *ATLAS* (task=call method=majorityBase) to produce haploid calls on known alleles (1240K sites; (Mathieson *et al*., 2015)) while taking PMD and recalibration parameters into account for the whole-genome samples. We merged our calls with the 1240K reference panel using a custom script and plink (v1.9; (Chang *et al*., 2015)) and converted them into *EIGENSTRAT* format (*convertf -p*, https://reich.hms.harvard.edu/software/InputFileFormats). We then used *hapROH* (v0.3a4; (Ringbauer, Novembre and Steinrücken, 2021)) with default parameters to identify runs of homozygosity (ROHs) in whole-genome modern and ancient samples. The *hapROH* output files were merged and the ROHs were binned on genetic length ranges (2-5cM, 5-10cM and >10cM) to calculate the total length and number of ROHs for each bin.

#### Phenotyping prediction

##### HIrisPlex analysis

We used *ATLAS* (taks=call method=MLE) to produce diploid calls on a set of 41 SNPs (Chaitanya *et al*., 2018) while taking PMD and recalibration parameters into account for our nuclear genomic data. We then used the HIrisPlex-S webtool (Walsh *et al*., 2014, 2017; Chaitanya *et al*., 2018) to predict possible pigmentation phenotypes. We therefore created a csv file by converting genotypes to allele counts for all alleles of interest needed for the prediction and uploaded it at https://hirisplex.erasmusmc.nl/. We obtained posterior probabilities for four phenotypes, each of them consisting of different categories that sum up to one: skin pigmentation (dark, intermediate, pale and very pale), hair pigmentation (black, brown, blond and red), eye pigmentation (brown, intermediate and blue) and hair shade (dark and light) (Figure S5). In order to reduce noise, we turned phenotypes with more than two categories into a binary form by summing up the corresponding probabilities falling in the darker (dark, intermediate, black and brown) or lighter (pale, very pale, blonde, red and blue) spectrum. We then proceeded to train a LDA model based on the summarized probabilities as input and using a set of Meso European-like and Neo Aegean-like samples as a grouping factor, excluding admixed samples and those from a different geographical context (see Figure 4).

## Supplemental Information

Document S1: Supporting Figures S1–S5 and Table S1.

Extended data tables:

Supplementary Data Table 1: Sample processing including detailed information on library preparation, read- and sample statistics, isotope-haplogroup and contamination information for sequenced libraries and ancient reference samples.

Supplementary Data Table 2: List of samples used in the interval-set and guidance set for local in-del realignment.

## Acknowledgements

This work was supported by Swiss National Science Foundation (31003A_173062 and 310030_200420 to DW, 31003A_182577 to MC), Marie Skłodowska-Curie actions ITN ‘‘BEAN’’. ZH was further supported by the European Research Council (856453 ERC-2019-SyG), Czech Grant Agency (GACR 21-17092X) and EMBO Long-Term Fellowship (ALTF 445-2017). CB was supported by the Fyssen Foundation (Fondation Fyssen, post-doctoral research grant).

We also thank the IBU cluster and sequencing facility of the University of Bern.

GENterprise, SAPM Munich,

## Document S1

**Figure S1.**
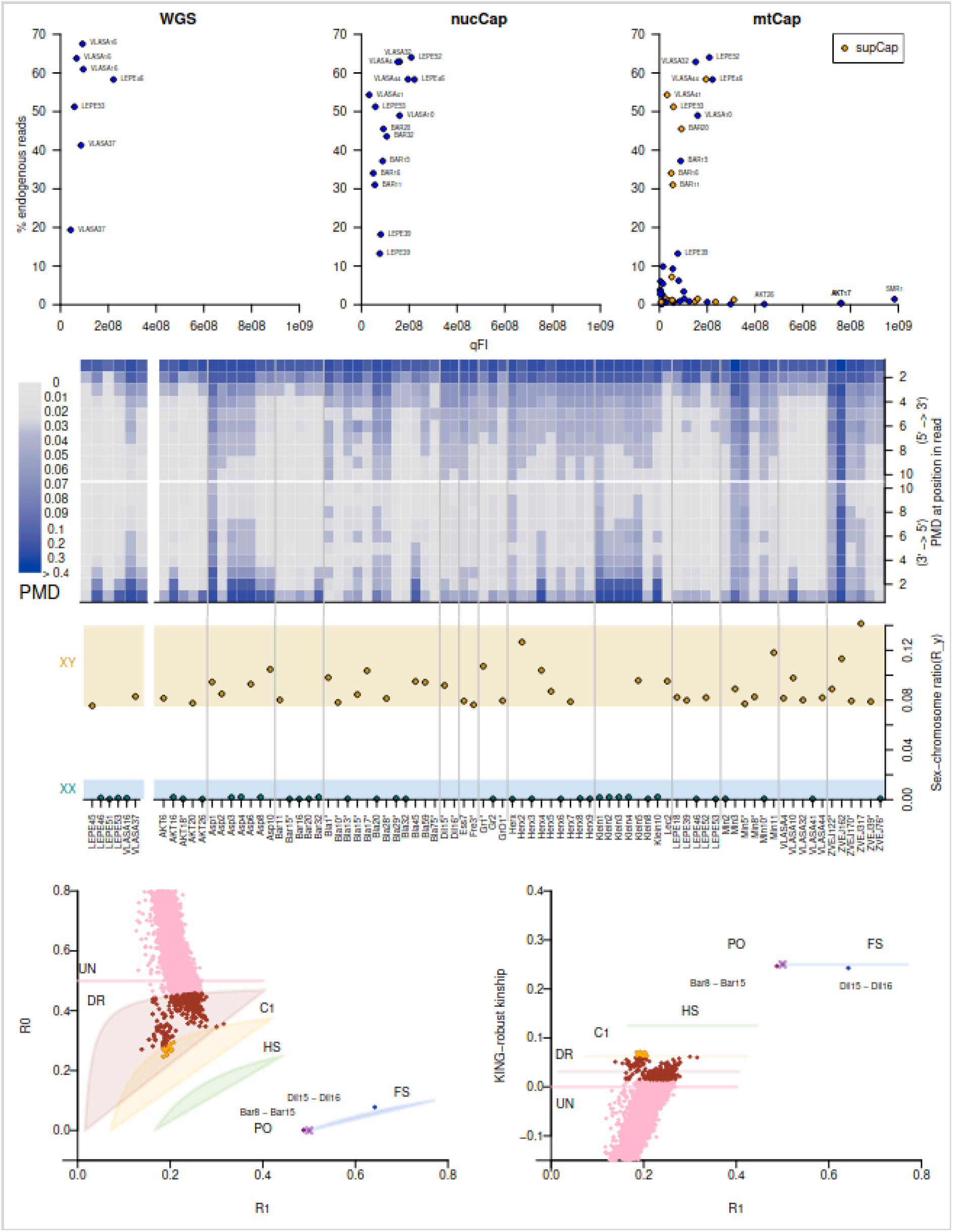
Quality assesment, sex and relatedness of the genomic samples. **(A)** Results from shallow MiSeq-screen of Libraries. The number of molecules in the fill-in product, measured with quantitative real-time PCR (qFI) and the percentage of reads aligning to the human reference (% endogenous). Colors and panels according to the strategy chosen. SupCap stands for mitochondrial capture on the supernatant of nuclear capture experiment, containing unhybridized DNA. **(B)** Post-mortem damage estimates for the first and last 10 bp of the reads. All samples show expected damage patterns **(C)** Fraction of Y- to (X+Y)-chromosomal reads. Up from a ratio above 0.075 (orange), an individual is considered to have a XY genotype, while a ratio below 0.016 (green) is considered to arise from a XX genotype. (Skoglund 2013). **(D)** Relatedness between the samples that passed quality filters. PO: parent-offspring, FS: full-siblings, HS: half-siblings/avuncular/grandparent-grandchild, C1: first cousin, UR: unrelated, DR: distantly related. All samples are not more than distantly related except for Bar8 and Bar15 that are PO, and Dil15 and Dil16 that are FS. Bar15 and Dil15 were additionally filtered out. **Related to STAR Methods, Library preparation and Target enrichment, Ancient DNA Authenticity, Molecular Sex Determination and Kinship analysis.**

**Figure S2.**
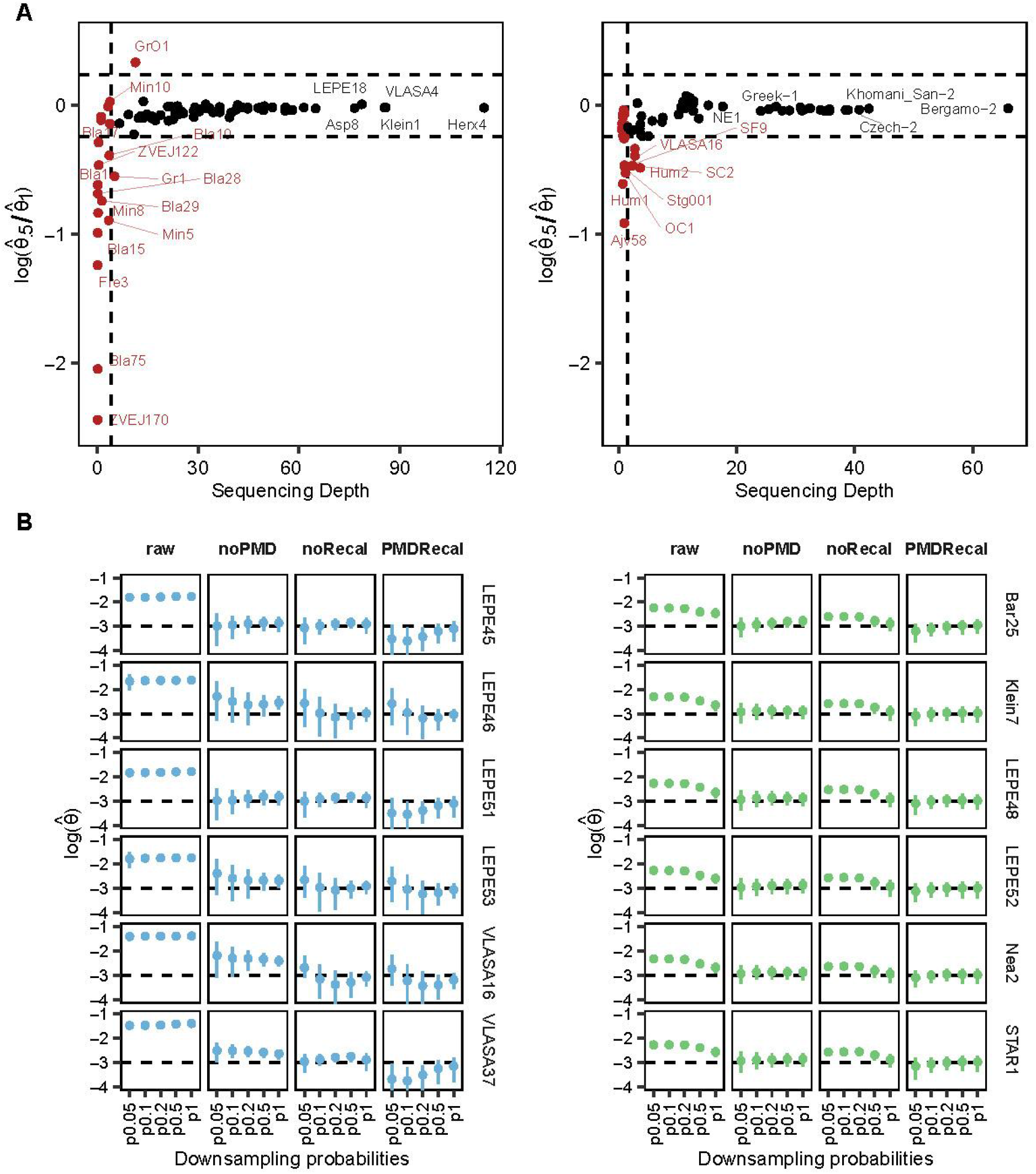
Downsampling analysis and quality filtering for comparability among samples. **(A)** log ratio between the median θ (heterozygosity) sampled estimates using all data and the median θ sampled estimates with data downsampled 50%. Additional quality filters were applied on all samples (left for capture, right for whole-genomes) based on the log ratio and sequencing depth (>1.5x and >4x for whole-genome and capture, respectively); both denoted as horizontal and vertical dotted lines. Samples coloured in red were discarded for downstream analysis. **(B)** θ downsampling experiments in low (1-4x, left) and medium-coverage (∼10x, right) samples from 100% to 5% data. Four different tests were done. Raw, the θ window estimates were calculated without taking into account post-mortem damage (PMD) and recalibration of base quality scores; noPMD, just recalibration parameters were considered; noRecal, only PMD parameters were used; PMDRecal,; both PMD and recalibration estimates were taken into account, this is the final way all our data were processed. **Related to STAR Methods, Depth and Heterozygosity Filtering, and Downsampling experiments.**

**Figure S3.**
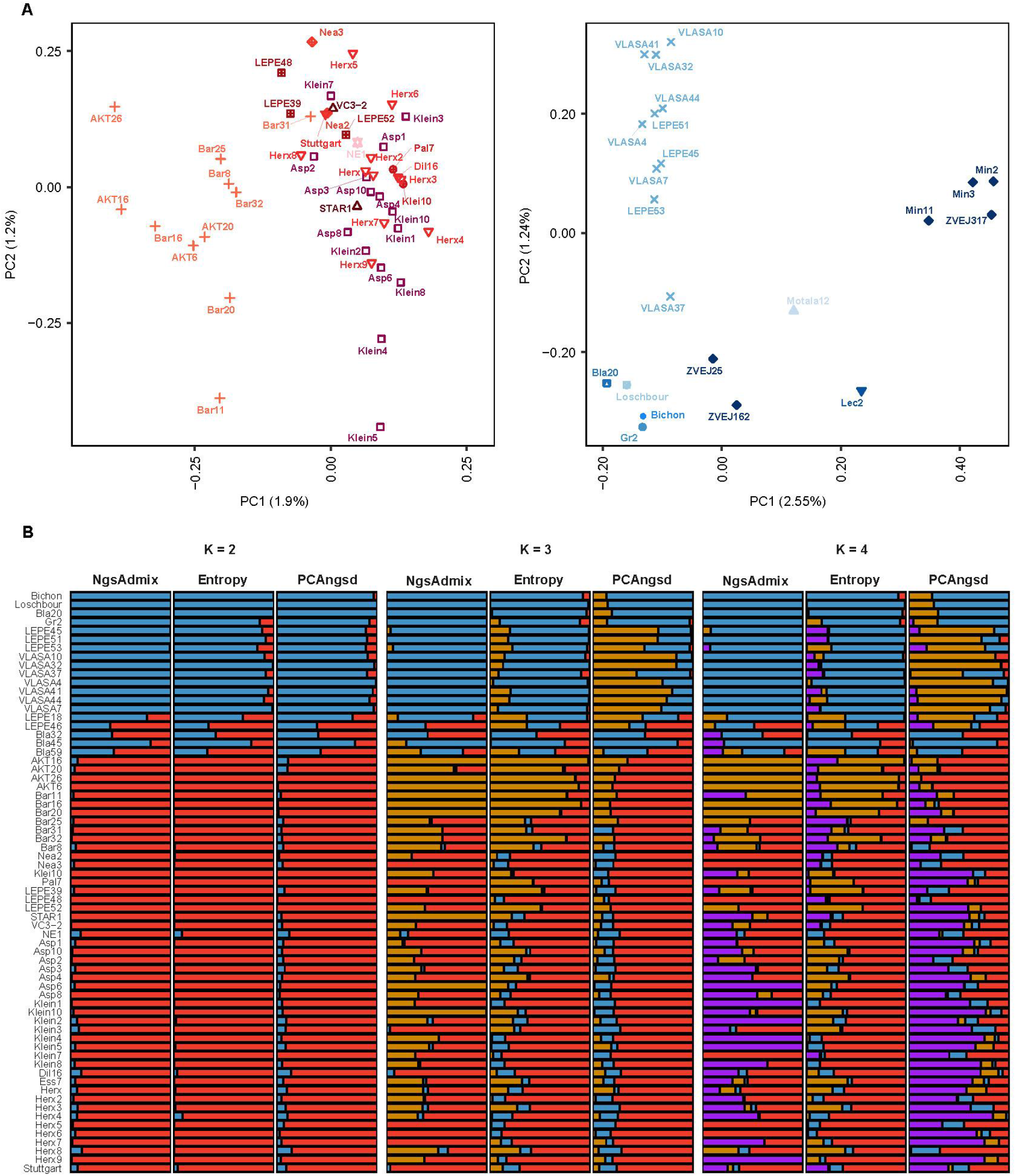
Genetic structure in the ancient samples. **(A)** PCAs for Neo Aegean-like and Meso European-like with their respective sample labels. **(B)** Admixture results from K=2 to K=4 using different softwares. The Evanno method and the deviance information criterion (DIC) were used for NGsAdmix and Entropy, respectively, to estimate the best K (PCAngsd estimates the best K on the fly). In all cases K=2 is the best. The set of samples used here excludes Lec2, Motala12 and North-Eastern Europeans compared to PCA. **Related to** Figure 2 **and STAR Methods, PCA and Admixture analysis.**

**Figure S4.**
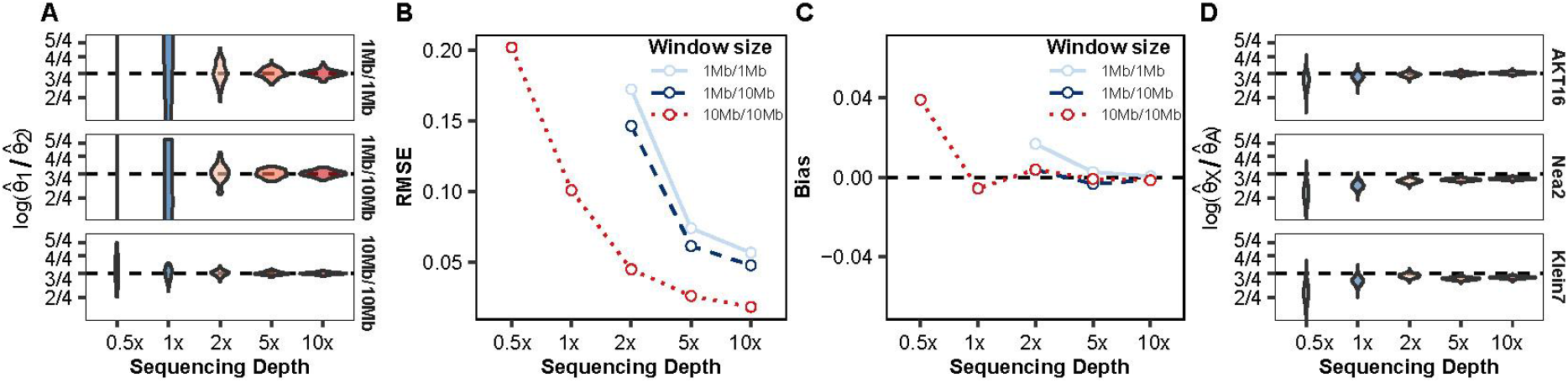
Theta ratio power analysis. **(A)** 100 BAMfiles were simulated with a sequencing depth of 10x and two chromosomes of same or different length (1Mb, 10Mb). The first chromosome was simulated with θ = 0.75*0.001 and the second with θ = 0.001 to mimic the expected ratio between the X chromosome and an autosome in humans. All BAMfiles were downsampled from 10x to 0.5x. The distribution of posterior medians obtained with ATLAS task=thetaRatio; at 0.5x most medians were out the plotted range. **(B)** Root mean squared error (RSME) between the posterior median and true value log(¾); when less than 10Mb are available per region below 2x, the RMSE becomes much larger than the scale of the plot. **(C)** The difference between the median of each 100 posterior medians per sequencing depth and window size and the true value log(¾); below 2x and with less than 10Mb per region the Bias becomes more negative than the scale of the plot. **(D)** Three ancient samples with medium coverage depth were chosen and downsampled from 10x to 0.5x to estimate the theta ratio between neutral sites in X chromosome and autosomes. Expected values are in dashed lines. **Related to** Figure 3 **and STAR Methods, Heterozygosity ratio**.

**Figure S5.**
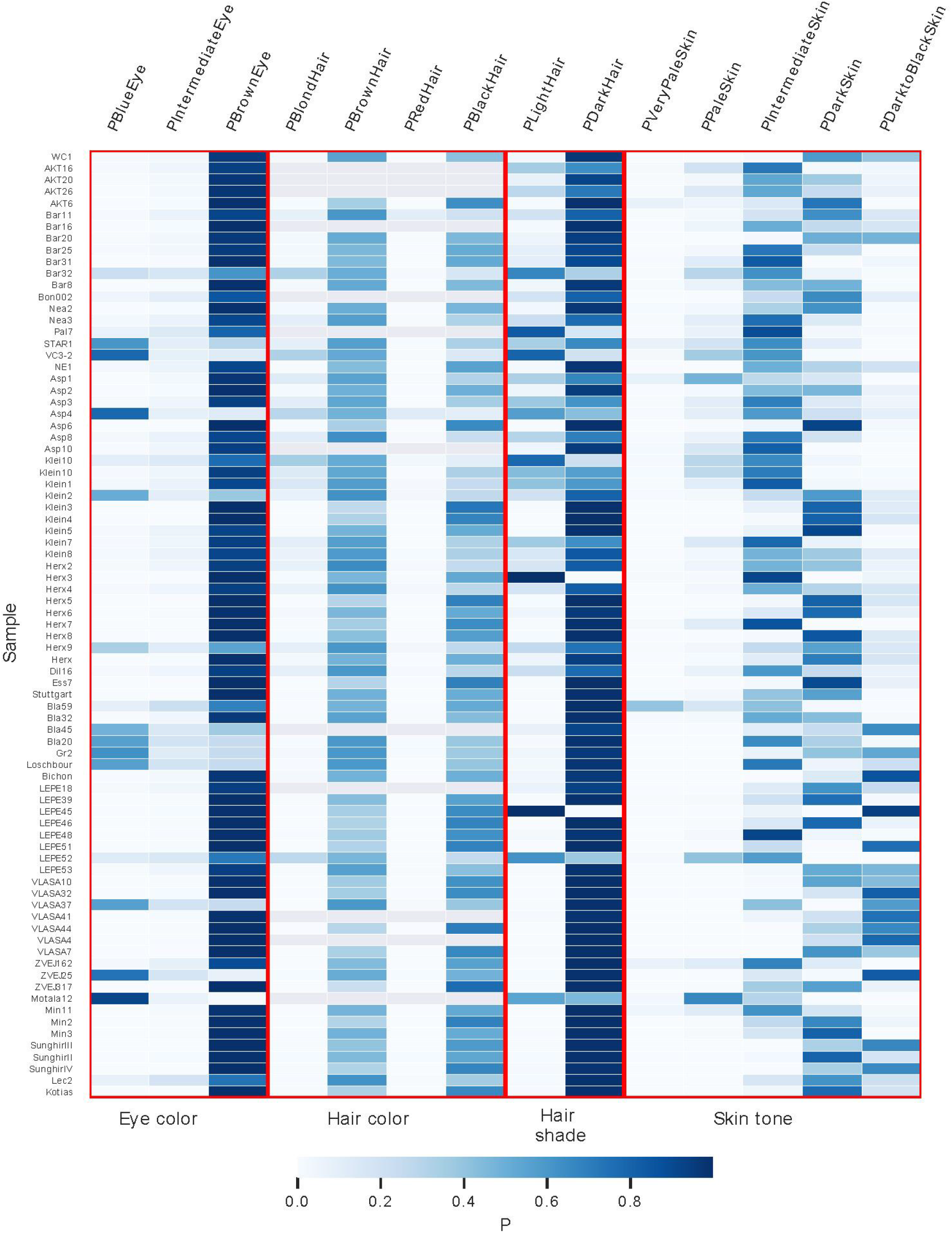
HIrisPlex analysis. Probabilities for eye, hair and skin pigmentation phenotypes estimated using the HIrisPlex-S webtool for newly sequenced capture and whole genomes and previously published individuals. In case an individual has capture and whole-genome sequences available, the analysis with capture data was just shown. Some SNPs associated with hair colour genotypes could not be called properly for some individuals. **Related to** Figure 4 **and STAR Methods, HIrisPlex analysis**

**Table S1.** Reference whole-genome samples.

| Sample ID | Site | Phase | Archeological/<br>BurialID | Age (cal. BC) | Mean<br>Depth | Sex | Publication |
| --- | --- | --- | --- | --- | --- | --- | --- |
| SunghirII | Sunghir | MUP | 2, SII | 32,234 ± 1049 | 3.86 | XY | (Sikora et al. 2017) |
| SunghirIII | Sunghir | MUP | 2, SIII | 32,092 ± 1061 | 10.16 | XY | (Sikora et al. 2017) |
| SunghirIV | Sunghir | MUP | 2, SIV | 31,992 ± 493 | 3.68 | XY | (Sikora et al. 2017) |
| SunghirI* | Sunghir | MUP | 1, SI | 30,822 ± 1052 | 1.05 | XY | (Sikora et al. 2017) |
| Kotias | Kotias Klde | LM | Layer A2 | 7,712 ± 183 | 11.93 | XY | (Jones et al. 2015) |
| ZVEJ25 | Zvejnieki | LM (Narva) | 93 | 5,738 ± 102.5 | 3.16 | XY | (Jones et al. 2017) |
| ZVEJ32* | Zvejnieki | MM (Kunda) | 313 | 6,358 ± 109 | 0.98 | XX | (Jones et al. 2017) |
| Hum1* | Hummervikholmen | M Scand. | - | 7,363.5 ± 88.5 | 0.7 | XX | (Günther et al. 2018) |
| Hum2* | Hummervikholmen | M Scand. | - | 7,363.5 ± 88.5 | 3.64 | XY | (Günther et al. 2018) |
| SF9* | Stora Förvar | M Scand. | G/Layer 9 | 7,144 ± 156 | 1.03 | XX | (Günther et al. 2018) |
| Stg001* | Steigen | M Scand. | - | 3,857 ± 93 | 1.11 | XY | (Günther et al. 2018) |
| Motala12 | Östergötland | M Scand. | - | 5,938.5 ± 422.5 | 1.92 | XX | (Lazaridis et al. 2014) |
| Satsurbia* | Satsurbia | UP | Area B, Y5 | 11,256 ± 124 | 1.17 | XY | (Jones et al. 2015) |
| Bichon | Grotte du Bichon | UP | Bichon | 11,665 ± 105 | 7.47 | XY | (Jones et al. 2015) |
| Loschbour | Heffingen | LM | LBK380 | 6,105 ± 115 | 12.05 | XY | (Lazaridis et al. 2014) |
| LaBraná* | La Braña-Arintero | M | La Braña 1 | 5,815 ± 125 | 2.71 | XY | (Olalde et al. 2014) |
| Canes1-Meso* | Canes | M | I-A | 5,115 ± 130 | 0.49 | XX | (González-Fortes et al. 2017) |
| Chan-Meso* | Chan do Lindeiro | M | Elba | 7,131 ± 124 | 0.95 | XX | (González-Fortes et al. 2017) |
| OC1-Meso* | Ostrovul Corbului | M | 24 | 6,704 ± 269 | 1.2 | XY | (González-Fortes et al. 2017) |
| SC2-Meso* | Schela Cladovei | M | M96/3 | - | 2.341 | XY | (González-Fortes et al. 2017) |
| VLASA7 | Vlasac | LM | VL 31 | 6,552 ± 212 | 15.21 | XY | (Marchi et al. 2022) |
| VLASA32<sup>#</sup> | Vlasac | LM | VL 16 | 7,604.5 ± 136.5 | 12.65 | XY | (Marchi et al. 2022) |
| WC1 | Wezmeh Cave | EN | n-10 | 7,268.5 ± 186.5 | 12.72 | XY | (Broushaki et al. 2016) |
| AH1* | Tepe Abdul Hosein | EN | 13030 | - | 1.1 | XX | (Broushaki et al. 2016) |
| AH2* | Tepe Abdul Hosein | EN | 19001 | 7,980.5 ± 224.5 | 0.6 | XY | (Broushaki et al. 2016) |
| AH4* | Tepe Abdul Hosein | EN | 10035 | 7,979.5 ± 224.5 | 0.82 | XX | (Broushaki et al. 2016) |
| Bon002 | Boncuklu | N | ZHB; Grave 9 | 8,128 ± 151 | 5.67 | XX | (Kilinc et al. 2016) |
| AKT16<sup>#</sup> | Aktopraklik | EN | 89 D 14.1 | 6,547.5 ± 87.5 | 10.85 | XX | (Marchi et al. 2022) |
| Bar31 | Barcin | EN | L11W-546 | 6,328 ± 91 | 3.56 | XY | (Hofmanová et al. 2016) |
| Bar8<sup>1</sup> | Barcin | EN | M10/106 | 6,121 ± 91 | 7.38 | XX | (Hofmanová et al. 2016) |
| Bar25 | Barcin | EN | BH 43347,<br>M10, 455 (lot.<br>1856) | 6,294.5 ± 89.5 | 12.66 | XY | (Marchi et al. 2022) |
| Nea3 | Nea Nikomedeia | EN | T XII | 6,183.5 ± 143.5 | 11.57 | XX | (Marchi et al. 2022) |
| Nea2 | Nea Nikomedeia | EN | #7 | 6,098 ± 75 | 12.51 | XX | (Marchi et al. 2022) |
| Rev5* | Revenia | EN | Rev5, burial 2 | 6,351 ± 87 | 1.14 | XX | (Hofmanová et al. 2016) |
| Klei10 | Kleitos | FN | grave 9 | 4,116 ± 118 | 2.49 | XY | (Hofmanová et al. 2016) |
| Pal7 | Paliambela | LN | Pal7 | 4,401 ± 51 | 1.56 | XX | (Hofmanová et al. 2016) |
| LEPE52<sup>#</sup> | Lepenski-Vir | E-MN | LV 73 | 5,812 ± 119 | 12.38 | XY | (Marchi et al. 2022) |
| LEPE48 | Lepenski-Vir | TEN | LV 122 | 5,939.5 ± 72.5 | 10.93 | XY | (Marchi et al. 2022) |
| STAR1 | Grad-Starčevo | EN<br>(Starčevo) | grave 1 | 5,532.5 ± 56.5 | 10.56 | XX | (Marchi et al. 2022) |
| VC3-2 | Vinča-Belo Brdo | EN<br>(Starčevo) | grave V - ND | 5,495.5 ± 69.5 | 11.23 | XY | (Marchi et al. 2022) |
| KO1* | Tiszaszőlös Domaháza | EN (Körös) | 4 | 5,715 ± 65 | 0.73 | XY | (Gamba et al. 2014) |
| NE1 | Polgár-Ferenci-hát | MN (ALP) | 325 V | 5,190 ± 120 | 17.66 | XX | (Gamba et al. 2014) |
| Klein7 | Kleinhadersdorf | EN (LBK) | Grave 56<br>(25.937) | 4,977 ± 181 | 11.3 | XX | (Marchi et al. 2022) |
| Asp6<sup>#</sup> | Asparn-Schletz | EN (LBK) | Ind 44, 646 Part<br>152, Schnitt 10<br>LM70 | 5,524.5 ± 50.5 | 12.12 | XY | (Marchi et al. 2022) |
| Dil16<sup>#,1</sup> | Dillingen-Steinheim | EN (LBK) | Grave 24,<br>Befund 24 | 5,116.5 ± 118.5 | 10.61 | XY | (Marchi et al. 2022) |
| Ess7<sup>#</sup> | Essenbach Ammerbreit | EN (LBK) | Grave 2 | 4,975 ± 75 | 12.34 | XY | (Marchi et al. 2022) |
| Herx<sup>#</sup> | Herxheim | EN (LBK) | 281-19-6 | 5,078 ± 85 | 11.47 | XX | (Marchi et al. 2022) |
| Stuttgart | Stuttgart Mühlhausen | EN (LBK) | grave (I-78,<br>area-1) | 4,900 ± 150 | 13.59 | XX | (Lazaridis et al. 2014) |
| CB13* | Cova Bonica | EN (Cardial) | CB13-HH34-IV 2 -2407 | 5,415 ± 55 | 0.84 | XX | (Olalde et al. 2015) |
| ZVEJ27* | Zvejnieki | M/EN | 121 | 5,077 ± 225 | 0.7 | XY | (Jones et al. 2017) |
| ZVEJ31* | Zvejnieki | MN (Ware) | 221 | 4,014 ± 214.5 | 1.01 | XX | (Jones et al. 2017) |
| GB1-Eneo* | Gura Baciului | Eneolithic | M1 | 3,377 ± 77 | 2.73 | XX | (González-Fortes et al.<br>2017) |
| Ajv58* | Ajvide | N (PWC) | - | 2,750 ± 150 | 0.96 | XY | (Skoglund et al. 2014) |
| KhSan-1 | South Africa | Modern | - | - | 38.45 | XX | (Mallick et al. 2016) |
| KhSan-2 | South Africa | Modern | - | - | 42.42 | XX | (Mallick et al. 2016) |
| Mbuti-2 | Congo | Modern | - | - | 30.81 | XX | (Mallick et al. 2016) |
| Mandenka-2 | Senegal | Modern | - | - | 32.44 | XX | (Mallick et al. 2016) |
| BaKenya-2 | Kenya | Modern | - | - | 31.34 | XX | (Mallick et al. 2016) |
| Turkish-2 | Turkey | Modern | - | - | 31.05 | XX | (Mallick et al. 2016) |
| Georgian-1 | Georgia | Modern | - | - | 35.32 | XY | (Mallick et al. 2016) |
| Greek-1 | Greece | Modern | - | - | 25.45 | XY | (Mallick et al. 2016) |
| Bulgarian-1 | Bulgaria | Modern | - | - | 34.83 | XY | (Mallick et al. 2016) |
| Albanian-1 | Albania | Modern | - | - | 24.09 | XX | (Mallick et al. 2016) |
| Hungarian-1 | Hungary | Modern | - | - | 27.7 | XX | (Mallick et al. 2016) |
| Czech-2 | Czech Republic | Modern | - | - | 40.77 | XY | (Mallick et al. 2016) |
| Polish-1 | Poland | Modern | - | - | 38.84 | XY | (Mallick et al. 2016) |
| Saami-1 | Finland | Modern | - | - | 35.71 | XX | (Mallick et al. 2016) |
| Finnish-1 | Finland | Modern | - | - | 32.18 | XX | (Mallick et al. 2016) |
| English-1 | England | Modern | - | - | 34.21 | XY | (Mallick et al. 2016) |
| French-2 | France | Modern | - | - | 26.68 | XX | (Mallick et al. 2016) |
| Spanish-2 | Spain | Modern | - | - | 35.72 | XX | (Mallick et al. 2016) |
| Bergamo-2 | Italy | Modern | - | - | 65.99 | XX | (Mallick et al. 2016) |
| Sardinian-2 | Italy | Modern | - | - | 28.96 | XX | (Mallick et al. 2016) |

<sup>a</sup> UP, Upper Palaeolithic; MUP, Mid Upper Palaeolithic; M, Mesolithic; EM, Early Mesolithic; MM, Middle Mesolithic; LM, Late Mesolithic; TEN, Transformational/Early Neolithic; N, Neolithic; EN, Early Neolithic; E-MN, Early-Middle Neolithic; MN, Middle Neolithic; LN, Late Neolithic; FN, Final Neolithic
\* Samples were discarded for downstream analysis.
<sup>#</sup> Data for both whole genome and capture nuclear genome.
<sup>1</sup> Related samples, Bar8-Bar15; Dil15-Dil16.

